# Double-bond geometry determines fatty acid metabolic fate and ferroptosis sensitivity

**DOI:** 10.64898/2026.07.17.739092

**Authors:** Cynthia A. Harris, Shunji Kato, Yurika Otoki, Elisa Martin Perez, Dylan Boone, Juan Lorenzo B. Pablo, Anna Greka, James A. Olzmann

## Abstract

Ferroptosis is driven by the accumulation of oxidatively damaged membrane phospholipids, making membrane lipid composition a central determinant of cell death sensitivity. While fatty acid chain length and degree of unsaturation are well-established regulators of ferroptosis, whether fatty acid stereochemistry contributes to ferroptosis susceptibility is mostly unexplored. Here, we systematically screened structurally diverse fatty acids for their ability to modulate ferroptosis and unexpectedly identified trans-unsaturated fatty acids as potent sensitizers. Compared with its cis counterpart linoleic acid, the trans polyunsaturated fatty acid (PUFA) linoelaidic acid more strongly enhanced lipid peroxidation and promoted the accumulation of ferroptosis-susceptible phospholipid species. Unexpectedly, the trans monounsaturated fatty acid petroselaidic acid also sensitized cells to ferroptosis, whereas its cis stereoisomer petroselinic acid suppressed ferroptosis. Mechanistically, petroselaidic acid required stearoyl-CoA desaturase-dependent conversion to a PUFA, directly demonstrating that double-bond geometry can redirect fatty acid metabolic fate through altered recognition by lipid metabolic enzymes. Although linoelaidic acid and petroselaidic acid followed distinct metabolic pathways, both converged on phospholipid remodeling that expanded pools of ferroptosis-susceptible membrane lipids. Together, our findings demonstrate that fatty acid double-bond geometry determines their metabolic fate and the membrane phospholipid composition, establishing lipid stereochemistry as a previously unrecognized structural determinant of ferroptosis sensitivity.

## INTRODUCTION

Ferroptosis is an iron-dependent form of regulated cell death induced by the overwhelming buildup of oxidatively damaged membrane phospholipids^1–4^. Consequently, ferroptosis sensitivity is governed by the balance between three interconnected processes: the generation of lipid radicals, the antioxidant defense systems that suppress lipid peroxidation, and the abundance of oxidation-prone membrane phospholipids that serve as substrates for peroxidation^1–4^. While substantial progress has been made in understanding the mechanisms governing antioxidant defense systems, such as those mediated by the glutathione-dependent peroxidase GPX4^5–7^ and the lipid antioxidant recycling enzyme FSP1^8,9^, increasing evidence indicates that membrane phospholipid composition is a fundamental determinant of ferroptosis sensitivity.

The susceptibility of membrane phospholipids to peroxidation is dictated by the chemical properties of their constituent fatty acids^4,10^. Polyunsaturated fatty acids (PUFAs) are intrinsically prone to free radical-mediated peroxidation because bis-allylic methylene groups facilitate hydrogen abstraction and lipid radical formation, whereas saturated and monounsaturated fatty acids (MUFAs) are comparatively resistant to oxidation. Accordingly, PUFA-containing phospholipids serve as the principal substrates for lipid peroxidation during ferroptosis^11,12^, whereas enrichment of membranes with MUFAs suppresses ferroptosis in part by limiting the abundance of oxidation-prone phospholipids. Indeed, supplementation with certain exogenous MUFAs displaces PUFAs from membrane phospholipids and potently protects cells from ferroptosis^13,14^. Emerging evidence further suggests that specific phospholipid species, including arachidonate-containing phospholipids^11,12^ and diPUFA phospholipids^15^, are particularly effective substrates for lipid peroxidation and ferroptosis. Indeed, the ratio of MUFAs and PUFAs in phospholipids has emerged as a central determinant of cancer cell ferroptosis sensitivity and therapeutic response^14,16–19^, underscoring the biological importance of understanding how membrane lipid composition is established and remodeled. Thus, the contribution of an individual fatty acid to ferroptosis is determined by two fundamental properties, its intrinsic chemical susceptibility to peroxidation and its incorporation into specific phospholipids.

The metabolic fate of fatty acids is governed by an interconnected network of lipid metabolic pathways that regulate fatty acid uptake, activation, desaturation, elongation, and phospholipid biosynthesis and remodeling^20^. Central to this process is the Lands cycle, which remodels membrane phospholipid composition through coordinated deacylation and reacylation reactions^4,10,21^. The acyl-CoA synthetase ACSL4 has emerged as a ferroptosis-driving enzyme that preferentially activates long-chain PUFAs for incorporation into phospholipids^11,12,22,23^, whereas lipid remodeling pathways that increase MUFA phospholipid content suppress ferroptosis^13^. More broadly, phospholipid remodeling determines both the abundance and molecular identity of oxidation-prone phospholipid species within cellular membranes, thereby shaping ferroptosis susceptibility. Although the chemical principles governing lipid peroxidation are well established, considerably less is known about the structural features that determine the metabolic fate of fatty acids and their incorporation into oxidation-prone membrane phospholipids.

Current paradigms have focused primarily on fatty acid chain length and degree of unsaturation as determinants of both chemical reactivity and ferroptosis sensitivity^4,10^. Despite these advances, our understanding of how fatty acid structure governs metabolic fate and phospholipid remodeling remains incomplete^20^. To systematically define how fatty acid structure regulates ferroptosis, we performed a functional screen of structurally diverse fatty acids. Unexpectedly, this unbiased approach identified trans unsaturated fatty acids as potent sensitizers of ferroptotic cell death, including the trans MUFA petroselaidic acid, which contrasts with the well-established ferroptosis-suppressive effects of cis MUFAs. Mechanistically, we demonstrate that double-bond geometry (i.e., cis vs trans) influences the metabolic fate of fatty acids, causing structurally distinct trans fatty acids to converge on the formation of oxidation-prone phospholipid species through distinct metabolic pathways. These findings establish fatty acid double bond geometry as a previously unrecognized structural determinant of membrane phospholipid composition and ferroptosis susceptibility.

## RESULTS

### Functional fatty acid screen identifies trans-unsaturated fatty acids as ferroptosis sensitizers

To systematically define how fatty acid structure influences ferroptosis sensitivity, we implemented the Fatty Acid Library for Comprehensive ONtologies (FALCON) screening platform^24^ to evaluate structurally diverse free fatty acids for their ability to modulate ferroptosis. A library of 62 fatty acids spanning saturated, monounsaturated, and polyunsaturated species with varying chain lengths and double-bond geometries was complexed to fatty acid-free BSA and H460 lung adenocarcinoma and U-2 OS osteosarcoma cells were incubated with a range of fatty acid concentrations in the presence or absence of the GPX4 inhibitor RSL3 (**Figure 1A**). Cell death was monitored by live-cell imaging and quantified as the normalized area under the lethal fraction curve (AUC) (**Figure 1A**).

**Figure 1.**
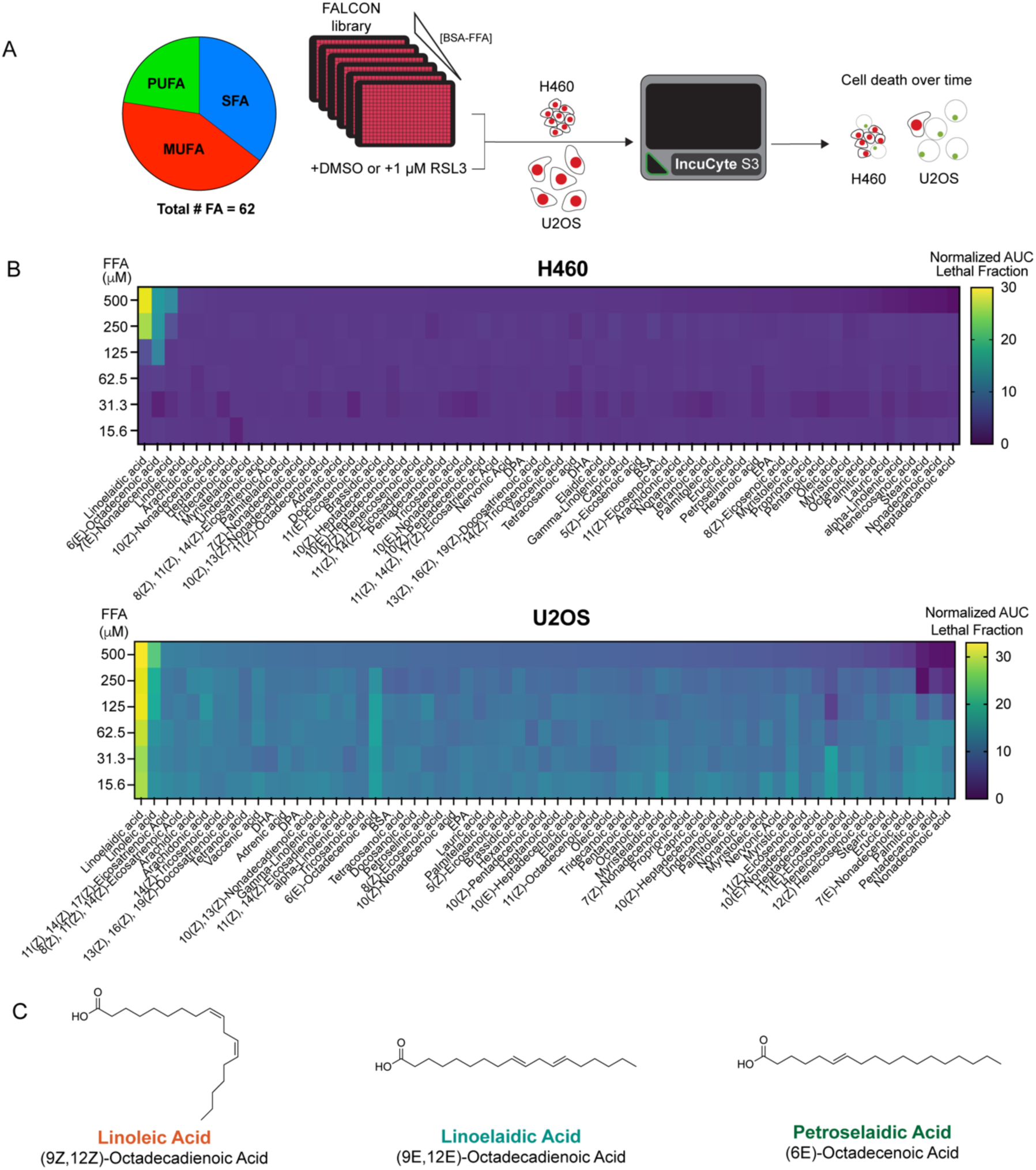
Functional lipid screening identifies trans-unsaturated fatty acids as potent ferroptosis sensitizers. (**A**) Outline of FFA functional screen in H460 and U-2 OS cells. (**B**) Normalized AUC of lethal fraction across the FALCON library in H460 (top) or U-2 OS (bottom). Normalization was conducted by subtracting the AUC of lethal fraction of the DMSO condition from the AUC of lethal fraction from the RSL3 condition. (**C**) Structures of hits from the FALCON screen with emphasized cis/trans stereochemistry.

The majority of fatty acids produced little or no enhancement of RSL3-induced cell death in either H460 or U-2 OS cells, consistent with ferroptosis sensitization not being a general property of exogenous fatty acid supplementation but rather sensitization depending on specific structural features of individual fatty acids (**Figure 1B**). Indeed, only a relatively small subset of fatty acids reproducibly enhanced ferroptotic cell death under these conditions. As expected based on previous studies, the cis PUFA linoleic acid (9Z,12Z)-Octadecadienoic Acid) enhanced ferroptosis^25^ (**Figure 1B,C**), supporting the validity of our screening approach.

Unexpectedly, the strongest hits identified were two structurally distinct trans-unsaturated C18 fatty acids (**Figure 1B,C**). The trans PUFA linoelaidic acid (9E,12E-octadecadienoic acid) ranked as the most potent sensitizer in both H460 and U-2 OS cells (**Figure 1B,C**). In addition, the trans MUFA petroselaidic acid (6E-octadecenoic acid) emerged as one of the strongest sensitizers despite containing only a single double bond (**Figure 1B,C**). In H460 cells, linoelaidic acid and petroselaidic acid represented the two highest-ranking sensitizing fatty acids identified in the screen, whereas in U-2 OS cells, linoelaidic acid was the strongest hit (**Figure 1B**). The identification of both a trans PUFA and a trans MUFA among the strongest sensitizers to GPX4 inhibition suggest that cis-trans double bond geometry may represent a structural determinant of ferroptosis susceptibility.

### Trans fatty acids sensitize cells to ferroptosis

To validate the screening results, we examined the ability of the top screening hits to sensitize H460 cells to GPX4 inhibition. Treatment with linoleic acid, linoelaidic acid, or petroselaidic acid alone had no effect on baseline H460 viability or growth rate (**Supplementary Figure S1A**). Co-treatment with linoelaidic acid, petroselaidic acid, or linoleic acid increased sensitivity to RSL3 in a concentration-dependent manner (**Figure 2A**). Consistent with the screening results, linoelaidic acid produced the greatest enhancement of ferroptotic cell death, whereas petroselaidic acid also robustly sensitized cells despite containing only a single double bond (**Figure 2A,B**).

**Figure 2.**
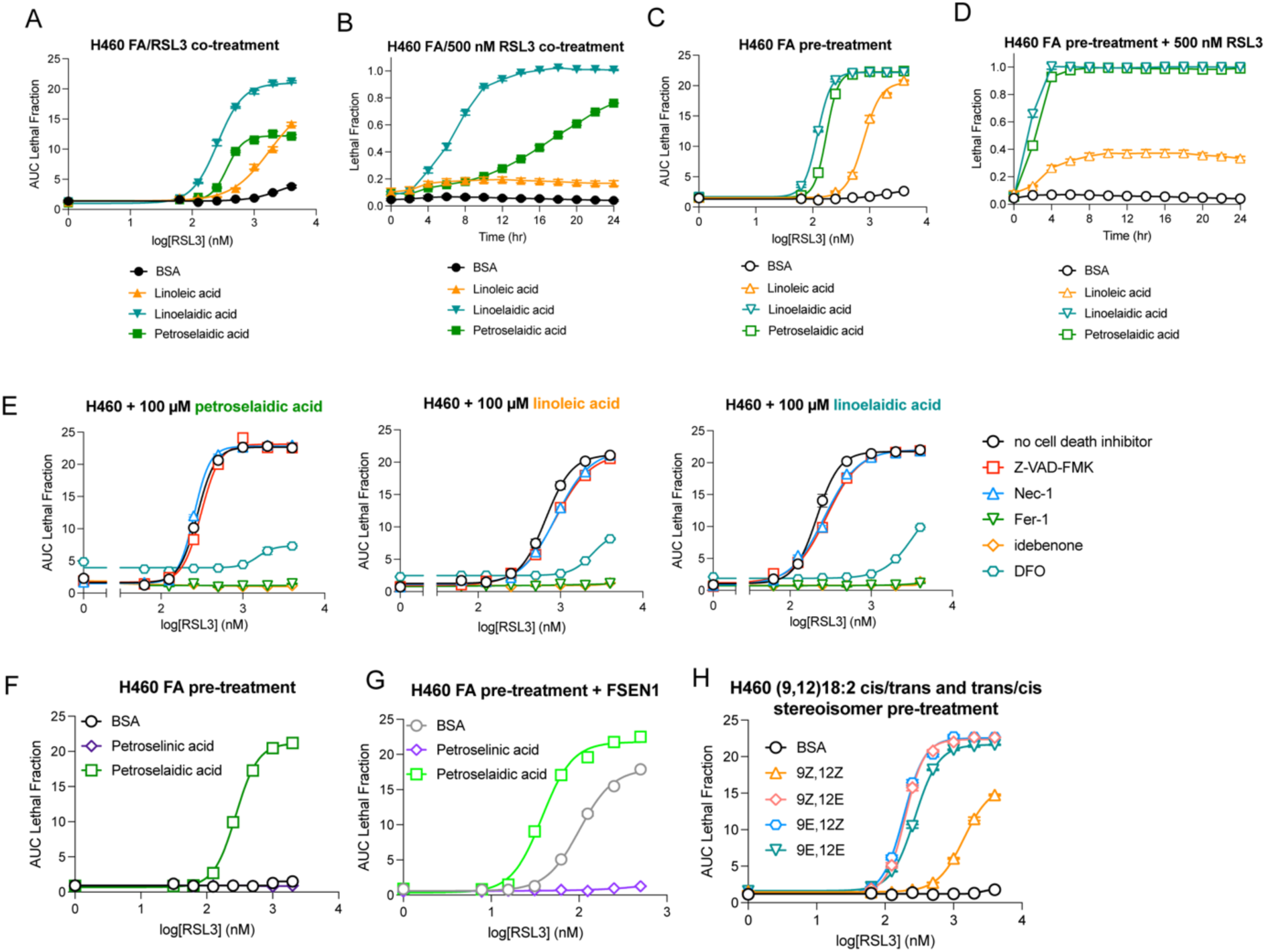
Trans fatty acids potently sensitize to ferroptosis. (**A**) RSL3 dose response of H460 cells co-treated with BSA or 100 µM indicated fatty acids. (**B**) Lethal fraction over time of H460 cells co-treated with 500 nM RSL3 and BSA or 100 µM indicated fatty acids. (**C**) RSL3 dose response of H460 cells pre-treated with BSA or 100 µM indicated fatty acids for 24 hours prior to RSL3 addition. (**D**) Lethal fraction over time of H460 cells pre-treated with BSA or 100 µM indicated fatty acids for 24 hours then treated with 500 nM RSL3. (**E**) Cell death rescue dose responses with RSL3 following 24 hour BSA or 100 µM fatty acid pre-treatment. Z-VAD-FMK, 20 µM; Necrostatin-1 (Nec-1), 1 µM; Ferrostatin-1 (Fer-1), 10 µM; idebenone, 10 µM; deferoxamine (DFO), 100 µM. (**F**) RSL3 dose response of H460 cells pre-treated with BSA or 100 µM indicated (6)-18:1 MUFA for 24 hours. (**G**) RSL3 dose response in the presence of 1 µM FSEN1 of H460 cells pre-treated with BSA or 100 µM indicated (6)-18:1 MUFA for 24 hours. (**H**) RSL3 dose response of H460 cells pre-treated with BSA or indicated (9,12)-18:2 PUFA for 24 hours.

Because exogenous fatty acids require cellular uptake, metabolic processing, and incorporation into membrane phospholipids before influencing membrane composition, we tested whether prolonged fatty acid exposure further enhanced ferroptosis sensitivity. Cells were pretreated with fatty acids prior to RSL3 exposure to allow additional time for membrane remodeling. Consistent with this hypothesis, pretreatment enhanced sensitization by both linoelaidic acid and petroselaidic acid compared with co-treatment, producing a pronounced leftward shift in the RSL3 dose-response curve and accelerating the kinetics of cell death (**Figure 2C,D**). In contrast, pretreatment with the cis PUFA linoleic acid produced only modest enhancement under identical conditions (**Figure 2C,D**). U-2 OS cells pre-treated with petroselaidic acid were also slightly sensitized to RSL3, though petroselaidic acid treatment itself was sufficient to induce cell death in a fraction of cells (**Supplementary Figure S1B**). Remarkably, pre-treatment of U-2 OS and HEK 293 cells with linoelaidic acid contributed to more of an increase in RSL3 sensitivity than both arachidonic acid and linoleic acid pre-treatments (**Supplementary Figure S1C,D**). This suggests that in certain cellular contexts, changes in ferroptosis sensitivity upon exogenous PUFA addition cannot be predicted by the degree of PUFA unsaturation alone.

Linoelaidic acid and petroselaidic acid also sensitized cells to the chemically distinct GPX4 inhibitors ML162 and ML210 (**Supplementary Figure S1E,F**), indicating that their effects are not specific to RSL3. Linoleic acid and linoelaidic acid pre-treatment sensitized cells to inhibition of glutathione synthesis upstream of GPX4 by the system Xc- inhibitors erastin-2 and imidazole ketone erastin (IKE) as well as the γ-glutamylcysteine synthetase inhibitor buthionine sulfoximine (BSO) (**Supplementary Figure S1G-I**). Interestingly, fatty acid pre-treatment did not enhance the toxicity of FINO_2_, an endoperoxide-containing compound that promotes iron oxidation in cells^26^, but fatty acid pre-treatment did subtly increase cell death in combination with FIN56, which has multi-faceted functions in ferroptosis induction^27^ (**Supplementary Figure S1J,K**).

By contrast, neither fatty acid promoted cell death when combined with the FSP1 inhibitor FSEN1 alone in wild-type H460 cells (**Supplementary Figure S2A**). H460 FSP1 knockout cells were similarly resistant to fatty acid treatment (**Supplementary Figure S2B**), consistent with GPX4 serving as the dominant ferroptosis defense pathway in this cell line. However, in GPX4-deficient H460 cells, both linoelaidic acid and petroselaidic acid synergized with FSP1 inhibition (**Supplementary Figure S2C**). Similar sensitization by both linoelaidic acid and petroselaidic acid was also observed in HT1080 fibrosarcoma cells, including a GPX4-deficient and FSP1-dependent genetic model (**Supplementary Figure S2D,E**). In GPX4-deficient, ferrostatin-1-dependent HT1080 cells, pre-treatment with linoelaidic acid followed by ferrostatin-1 washout enhanced cell death compared to BSA treatment (**Supplementary Figure S2F,G**). Indeed, pre-treatment with linoelaidic acid sensitized a broad panel of cell lines to RSL3, indicating that the ferroptosis-enhancing activity of trans fatty acids is not restricted to H460 and U-2 OS cells (**Supplementary Figure S2H-J**).

To determine whether the observed cell death is ferroptosis or another type of regulated cell death, we incubated cells with inhibitors of different cell death pathways. Cell death in the presence of RSL3 with linoelaidic acid and petroselaidic acid was strongly suppressed by incubation with ferroptosis inhibitors, including the lipophilic radical trapping antioxidants ferrostatin-1 and idebenone and the iron chelator deferoxamine (**Figure 2E**). In contrast, inhibitors of apoptosis (i.e., Z-VAD-FMK) or necroptosis (i.e., necrostatin-1) had little effect (**Figure 2E**). These findings demonstrate that trans fatty acid treatment enhances ferroptotic cell death.

The ferroptosis-promoting activity of petroselaidic acid was unexpected because exogenous cis MUFAs are widely recognized to suppress ferroptosis^13,14^. To directly examine the influence of double-bond geometry within MUFAs, we compared petroselaidic acid (6E-18:1) with its cis stereoisomer, petroselinic acid (6Z-18:1) (**Supplementary Figure S3A**). Whereas petroselaidic acid strongly sensitized H460 cells to RSL3, petroselinic acid exhibited little effect (**Figure 2F**). Because H460 cells express high levels of FSP1 and are highly resistant to ferroptosis^8^, any protective effect of cis MUFAs is masked by the already robust endogenous antioxidant defense. Indeed, pharmacological inhibition of FSP1 with FSEN1 sensitized H460 cells to ferroptosis (**Supplementary Figure S3B**) and unmasked a novel protective activity of the cis-MUFA petroselinic acid, while the trans MUFA petroselaidic acid continued to enhance ferroptosis (**Figure 2G**). Thus, cis and trans PUFAs exert strikingly opposite effects on ferroptosis despite differing only in their double-bond geometry.

To further examine the contribution of double-bond stereochemistry, we compared all four stereoisomers of (9,12)-18:2 fatty acids (**Supplementary Figure S3C**). Introduction of a single trans double bond was sufficient to enhance ferroptosis sensitivity, with the (9Z,12E), (9E,12Z), and (9E,12E) stereoisomers all producing substantially greater sensitization than the fully cis (9Z,12Z) isomer (**Figure 2H**). In contrast, the presence of two trans double bonds conferred little additional activity relative to the mixed cis/trans isomers, indicating that the acquisition of a trans double bond in an 18:2 fatty acid, rather than the total number of trans bonds, may be the dominant structural feature governing ferroptosis sensitization. Together, these studies validate the screening results and demonstrate that trans-unsaturated fatty acids promote ferroptosis through a mechanism that depends on double-bond geometry.

### Petroselaidic acid is metabolically converted into a trans PUFA through SCD-dependent desaturation

The finding that cis and trans MUFAs exert opposite effects on ferroptosis despite differing only in double-bond geometry suggests that stereochemistry fundamentally alters fatty acid metabolism. We therefore sought to determine the metabolic fates of linoleic acid, linoelaidic acid, or petroselaidic acid by performing temporal fatty acid profiling following treatment (**Figure 3A**). As expected, treatment with each fatty acid resulted in a rapid increase in the corresponding exogenous fatty acid species (**Figure 3A**). Linoleic acid treatment selectively increased cellular 18:2 fatty acids, whereas linoelaidic acid treatment increased trans 18:2 fatty acid abundance with minimal accumulation of other species (**Figure 3A**). Notably, though linoleic acid can be converted to more highly desaturated omega-6 PUFAs via elongation and desaturation, linoleic acid treatment only moderately increased the amount of gamma-linolenic acid (18:3n6) and did not significantly increase the amount of arachidonic acid (20:4n6) in these H460 cells over 24 hours, suggesting that linoleic acid’s ferroptosis-enhancing effects are not dependent on conversion to a more highly unsaturated PUFA (**Supplementary Figure S4A,B**). Inhibition of FADS2, the rate-limiting desaturase in the conversion of linoleic acid to gamma-linolenic acid en route to arachidonic acid, had no effect on RSL3 dose response in BSA, linoleic acid, linoelaidic acid, or petroselaidic acid pre-treated H460 cells (**Supplementary Figure S4C**).

**Figure 3.**
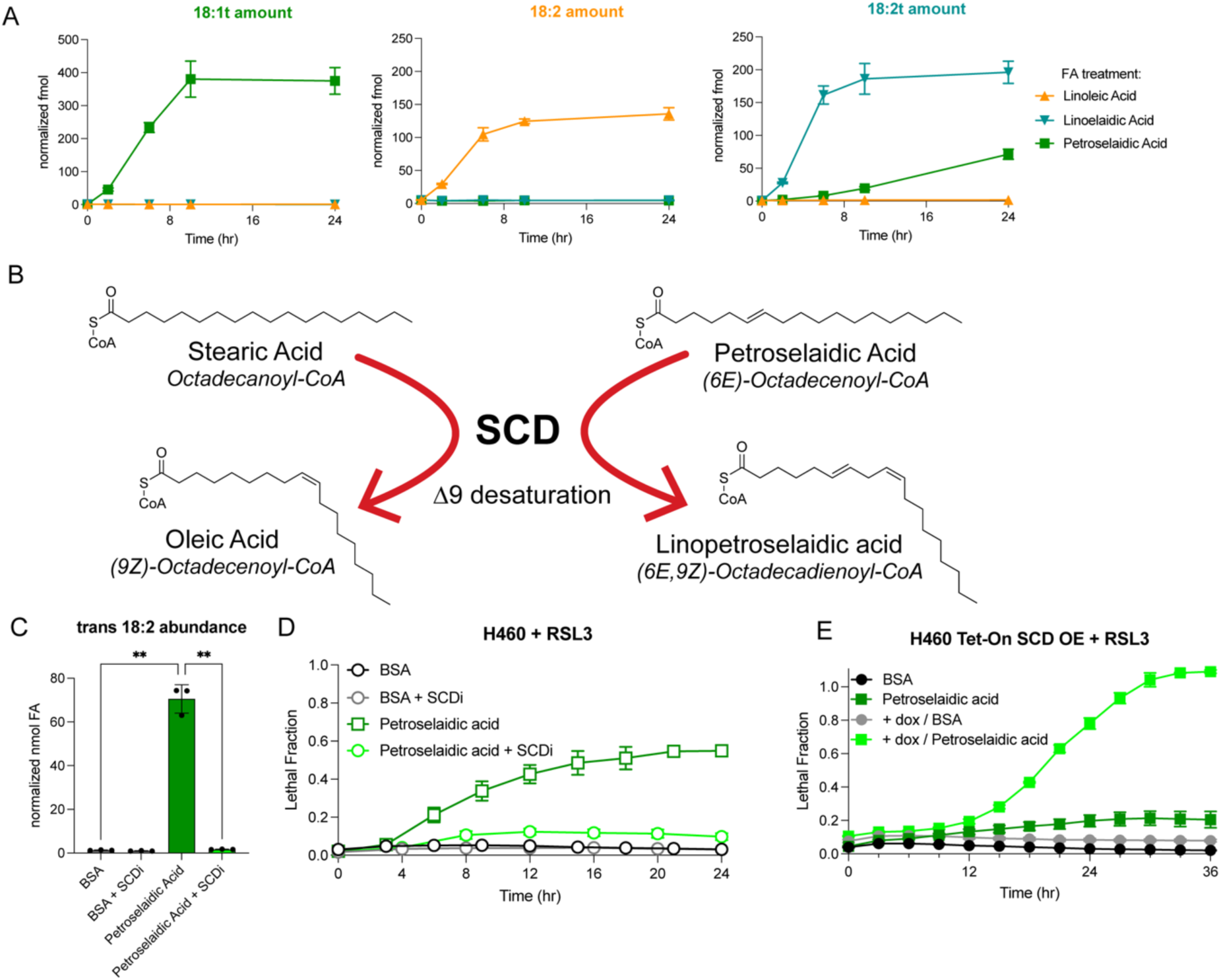
SCD-dependent desaturation redirects the metabolic fate of petroselaidic acid to promote ferroptosis. (**A**) GC-FID quantification of trans 18:1, 18:2, and trans 18:2 in H460 cells treated with 100 µM indicated fatty acid at time t = 0 hr. (**B**) Structures of native SCD substrate stearoyl-CoA and product oleoyl-CoA compared to structures of petroselaidyl-CoA and putative Δ9-desaturation product linopetroselaidyl-CoA. (**C**) GC-FID quantification of trans 18:2 in H460 cells treated for 24 hours with BSA or 100 µM petroselaidic acid in the presence or absence of 1 µM of the SCD inhibitor CAY10566. (**D**) Lethal fraction over time following addition of 125 nM RSL3 for H460 cells pre-treated with BSA or 100 µM petroselaidic acid in the presence or absence of 1 µM of SCD inhibitor for 24 hours. (**E**) Lethal fraction over time for wildtype or SCD overexpressing (+ dox) H460 cells co-treated with 2 µM RSL3 and BSA or 100 µM petroselaidic acid.

Petroselaidic acid treatment produced an unexpected metabolic profile. Although intracellular trans 18:1 levels rapidly increased following treatment, prolonged incubation resulted in the appearance of a trans 18:2 fatty acid species that was absent from control cells (**Figure 3A**). This observation suggested that petroselaidic acid may undergo metabolic conversion to a trans-bond containing 18:2 PUFA species.

Mammalian cells express multiple lipid desaturases with distinct substrate specificities and positional preferences (**Supplementary Figure S4D**). DEGS1 and DEGS2 catalyze desaturation of sphingoid bases during ceramide biosynthesis rather than fatty acids^28,29^, whereas the physiological substrates and functions of FADS3 remain incompletely defined but suggest a role in Δ14 desaturation of sphingoid bases^30^. Among the established fatty acid desaturases, FADS2 primarily introduces a Δ6 double bond with minor activity toward the Δ4 and Δ8 positions^31,32^, though the Δ6 position is already occupied by the pre-existing trans double bond of petroselaidic acid. FADS1 functions predominantly in long-chain PUFA biosynthesis and is not expected to utilize an 18-carbon PUFA as a substrate^33,34^. SCD catalyzes cis Δ9 desaturation of C16-C18 fatty acyl-CoAs^35^. Because petroselaidic acid contains a trans double bond at Δ6, SCD-mediated Δ9 desaturation would be predicted to be possible and generate a trans/cis PUFA (6E,9Z)-18:2 linopetroselaidic acid (**Figure 3B**). Moreover, the relatively linear conformation of petroselaidic acid, similar to SCD’s native saturated fatty acyl substrate stearoyl-CoA^36^, raises the possibility that it could serve as an unconventional substrate for SCD despite already containing a double bond^37,38^.

To directly test this desaturase-driven model, cells were treated with petroselaidic acid in the presence of the SCD inhibitor CAY10566. Pharmacological inhibition of SCD reduced the level of expected MUFAs (e.g., oleate and palmitoleate) (**Supplementary Figure S4E,F**). Strikingly, SCD inhibitor treatment completely prevented the appearance of the trans-18:2 metabolite, demonstrating that formation of this species requires SCD activity (**Figure 3C**). SCD inhibition also abolished the ability of petroselaidic acid to sensitize cells to RSL3-induced ferroptosis, reducing cell death to levels observed in control cells (**Figure 3D**) and demonstrating that a portion of the ferroptosis promoting effects of petroselaidic acid require metabolic conversion. The ferroptosis-inhibiting effect of SCD inhibitor treatment in the presence of petroselaidic acid is notable because SCD inhibitor treatment promotes RSL3-induced ferroptosis in the absence of petroselaidic acid (**Supplementary Figure S4G,H**), likely through the depletion of anti-ferroptotic MUFAs.

If SCD-mediated conversion of petroselaidic acid generates the bioactive ferroptosis-sensitizing metabolite, then increasing SCD activity should further enhance this response. To test this prediction, we generated cells with doxycycline-inducible SCD overexpression. Induction of SCD expression potentiated petroselaidic acid-dependent ferroptosis following GPX4 inhibition, whereas either treatment alone had little effect (**Figure 3E, Supplementary Figure S4I**). Our complementary loss- and gain-of-function experiments demonstrate that SCD activity is both necessary and sufficient to promote the ferroptosis-sensitizing activity of petroselaidic acid.

Collectively, these findings reveal an unexpected metabolic pathway whereby the trans MUFA petroselaidic acid is converted by SCD into a PUFA. Rather than functioning as a conventional MUFA, petroselaidic acid undergoes enzymatic desaturation to generate a metabolite that promotes ferroptosis, establishing fatty acid stereochemistry as a determinant of SCD substrate utilization and metabolic fate.

### Trans fatty acid sensitization is independent of triacylglycerol synthesis and lipid droplet formation

Exogenous fatty acids may influence ferroptosis through multiple mechanisms, including direct effects as free fatty acids or following metabolic activation and incorporation into complex lipids. To distinguish between these possibilities, we examined whether metabolic activation was required for ferroptosis sensitization. Inhibition of long-chain acyl-CoA synthetases with triacsin C abolished ferroptosis sensitization by both linoelaidic acid and petroselaidic acid (**Supplementary Figure S5A,B**), demonstrating that metabolic activation is required for their activity.

Once activated, fatty acids have multiple potential metabolic fates, including incorporation into membrane phospholipids, storage as triacylglycerols (TAGs), or further remodeling through lipid metabolic pathways. Because sequestration into TAGs and lipid droplets limits fatty acid incorporation into membrane phospholipids and suppresses ferroptosis^39–45^, one possibility that we considered was that trans fatty acids evade neutral lipid storage, and thereby increase their availability for incorporation into oxidation-prone membrane phospholipids. To examine this possibility, H460 cells were treated with linoleic acid, linoelaidic acid, or petroselaidic acid and lipid droplets were visualized by BODIPY 493/503 staining. All three fatty acids robustly induced lipid droplet formation compared with BSA-treated controls (**Figure 4A**). Quantification revealed similar increases in lipid droplet abundance following treatment with each fatty acid (**Figure 4B; Supplementary Figure S5C,D**). Consistent with these observations, flow cytometric analysis demonstrated comparable increases in neutral lipid accumulation following treatment with each fatty acid (**Figure 4C**). Thus, the enhanced ferroptosis sensitization produced by the trans fatty acids is not simply due to a failure to induce neutral lipid storage.

**Figure 4.**
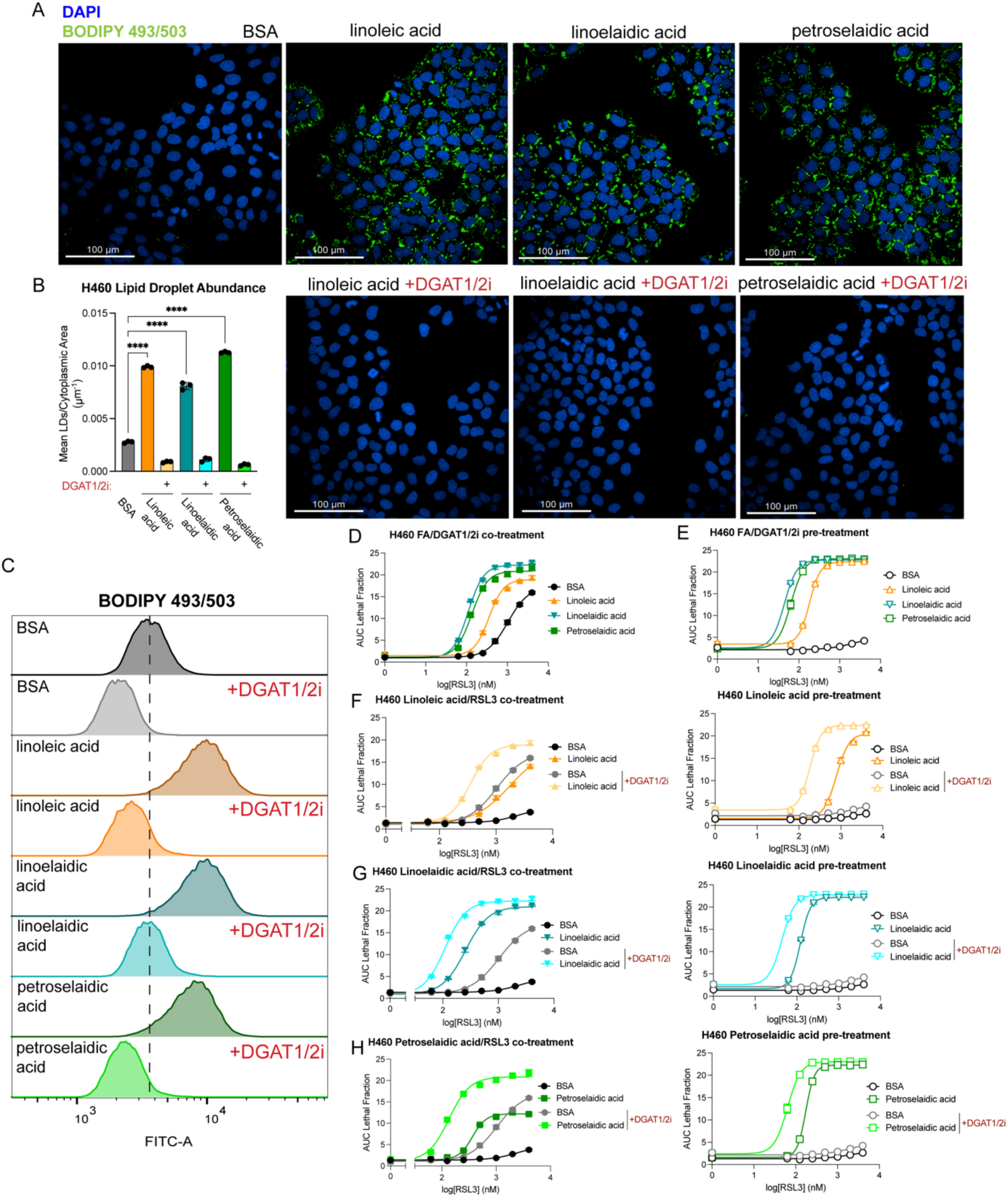
Trans fatty acid-mediated ferroptosis sensitization is independent of triacylglycerol synthesis and lipid droplet biogenesis. (**A**) Representative confocal fluorescence images of DAPI and BODIPY 493/503-stained H460 cells following treatment with BSA or 100 µM indicated fatty acid for 24 hours in the presence or absence of 45 µM A-922500 DGAT1 inhibitor and 30 µM PF-06424439 DGAT2 inhibitor. Scale bar represents 100 microns. (**B**) Quantification of mean number of lipid droplets per cytoplasmic area in H460 cells treated with the indicated fatty acids or BSA in the presence or absence of DGAT1 and DGAT2 inhibitors. (**C**) Flow cytometry analysis of BODIPY 493/503-stained H460 cells following treatment with BSA or 100 µM indicated fatty acid for 24 hours in the presence or absence of DGAT1 and DGAT2 inhibitors. The dashed vertical line represents the mean fluorescence intensity of BSA-treated cells. (**D**) RSL3 dose response of H460 cells co-treated with BSA or 100 µM indicated fatty acids and DGAT1 and DGAT2 inhibitors. (**E**) RSL3 dose response of H460 cells pre-treated with BSA or 100 µM indicated fatty acids and DGAT1 and DGAT2 inhibitors for 24 hours prior to RSL3 addition. (**F**) RSL3 dose response of H460 cells co-treated (left) or pre-treated (right) with BSA or 100 µM linoleic acid in the presence or absence or DGAT1 and DGAT2 inhibitors. (**G**) RSL3 dose response of H460 cells co-treated (left) or pre-treated (right) with BSA or 100 µM linoelaidic acid in the presence or absence or DGAT1 and DGAT2 inhibitors. (**H**) RSL3 dose response of H460 cells co-treated (left) or pre-treated (right) with BSA or 100 µM petroselaidic acid in the presence or absence or DGAT1 and DGAT2 inhibitors.

Pharmacological inhibition of DGAT1/2, the enzymes responsible for the final step of TAG synthesis, abolished lipid droplet formation induced by linoleic acid, linoelaidic acid, and petroselaidic acid as assessed by both fluorescence microscopy and flow cytometry (**Figure 4A- C**). Consistent with previous studies^39,40,42^, DGAT inhibition alone sensitized H460 cells to RSL3- induced ferroptosis (**Figure 4D,E**), supporting a protective role for TAG synthesis and lipid droplet formation. However, inhibition of DGAT1/2 failed to block ferroptosis sensitization by linoleic acid, linoelaidic acid, or petroselaidic acid under either co-treatment or pre-treatment conditions (**Figure 4D-H**). In U-2 OS cells, DGAT1/2 inhibition had minimal effect on RSL3 dose response under linoleic acid and linoelaidic acid pre-treatment (**Supplementary Figure S5E,F**). Interestingly, DGAT1/2 inhibition reversed the baseline of death seen with petroselaidic acid treatment alone in U-2 OS, though similarly to linoleic acid and linoelaidic acid pre-treatment in these cells, DGAT1/2 inhibition did not affect resistance to RSL3 (**Supplementary Figure S5G**). Arachidonic acid pre-treatment of U-2 OS in the presence of DGAT1/2 inhibitors resulted in slightly more resistance to RSL3 (**Supplementary Figure S5H**). These differences highlight the varied responses different cell lines employ in response to treatment with exogenous fatty acids.

Overall, these findings demonstrate that although trans fatty acids efficiently induce TAG synthesis and lipid droplet biogenesis, their ferroptosis-sensitizing activity is unrelated to altered neutral lipid storage.

### Trans PUFAs generate a distinct profile of lipid hydroperoxides during ferroptosis

Ferroptosis is ultimately characterized by the accumulation of oxidatively damaged membrane phospholipids. We therefore examined whether the enhanced ferroptosis sensitivity conferred by trans fatty acids is accompanied by increased membrane lipid peroxidation. H460 cells were pretreated with linoleic acid, linoelaidic acid, or petroselaidic acid prior to RSL3 treatment, and membrane oxidation was quantified using the lipid peroxidation probe BODIPY C11 and the lipid hydroperoxide probe Liperfluo. Consistent with its effects on ferroptosis sensitivity, linoelaidic acid pre-treatment produced substantially greater RSL3-induced membrane lipid peroxidation than linoleic acid, whereas petroselaidic acid produced a more modest increase (**Figure 5A, Supplementary Figure S5I**). Ferrostatin-1 suppressed lipid peroxidation under all conditions, confirming that the observed oxidation reflects ferroptotic lipid peroxidation (**Figure 5A**). Similar results were obtained using the lipid hydroperoxide-sensitive probe Liperfluo, which likewise demonstrated enhanced accumulation of lipid hydroperoxides following linoelaidic acid treatment, while petroselaidic acid failed to increase lipid hydroperoxide levels detected by this probe (**Figure 5B**). The more modest effects observed with petroselaidic acid contrasted with its robust ability to enhance ferroptosis, suggesting that oxidation of additional lipid species not detected by these bulk fluorescence assays may contribute to its activity.

**Figure 5.**
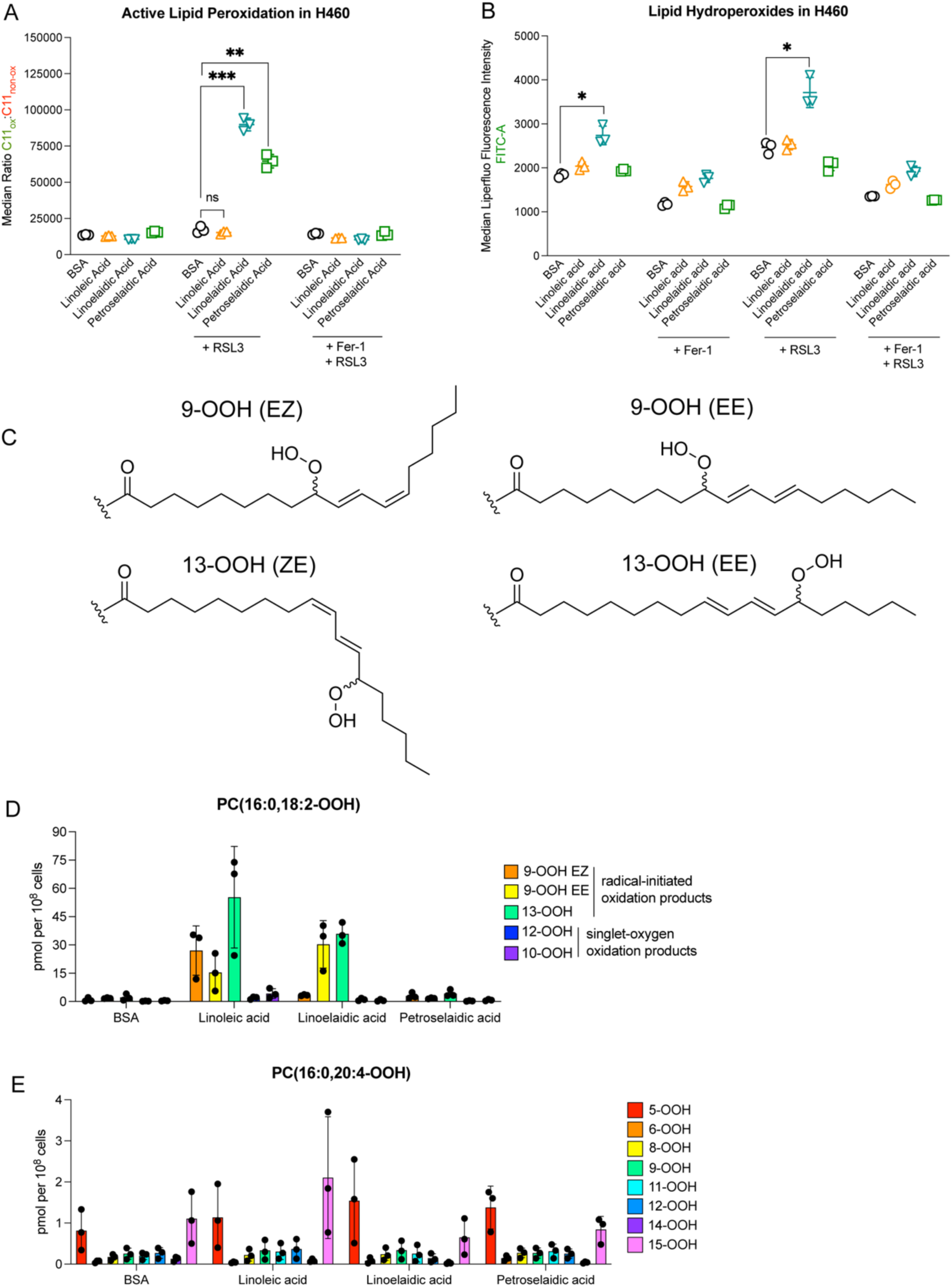
Double-bond geometry determines the identity of oxidized phospholipids during ferroptosis. (**A**) Median ratio of oxidized BODIPY C11 signal to non-oxidized BODIPY C11 signal in cells pre-treated with BSA or 100 µM indicated fatty acids for 20 hours followed by treatment with DMSO, 100 nM RSL3, or the combination of 100 nM RSL3 and 5 µM ferrostatin-1. (**B**) Median Liperfluo fluorescence intensity of cells pre-treated with BSA or 100 µM indicated fatty acids for 20 hours followed by treatment with DMSO, 5 µM ferrostatin-1, 100 nM RSL3, or the combination of 100 nM RSL3 and 5 µM ferrostatin-1. (**C**) Structures of hydroperoxyoctadecadienoyl (HpODE) products of radical initiated oxidation of (9,12)-18:2 species. (**D**) Quantification of different HpODE regiostereoisomers relative to PC(16:0,18:2-OOH) standards in H460 cells treated with BSA or 100 µM of the indicated fatty acids for 24 hours. 9- OOH EZ represents the trans-cis 9-HpODE, 9-OOH EE represents the trans-trans 9-HpODE, and 13-OOH represents the combined cis-trans and trans-trans 13-HpODEs. (**E**) Quantification of different arachidonate-derived hydroperoxyeicostetraenoyl (HpETE) regioisomers relative to PC(16:0,20:4-OOH) standards in H460 cells treated with BSA or 100 µM of the indicated fatty acids for 24 hours.

Radical-mediated oxidation of linoleate-containing phospholipids produces characteristic 9- and 13- hydroperoxide regioisomers, and the stereochemistry of the precursor fatty acid can influence the geometric configuration of these oxidation products^46^ (**Figure 5C**). Specifically, oxidation of cis linoleic acid-containing phospholipids is expected to generate mixed cis/trans hydroperoxide products, whereas oxidation of trans linoelaidic acid-containing phospholipids is expected to favor trans/trans hydroperoxide products^47^ (**Figure 5C**). To test this possibility, we performed targeted oxidized phospholipid lipidomics using phospholipid hydroperoxide standards to quantify individual oxidized phosphatidylcholine species. Consistent with oxidation of linoleate-containing phospholipids, linoleic acid treatment increased 9-OOH(EZ), 9-OOH(EE), and 13-OOH species (**Figure 5D**). In contrast, linoelaidic acid treatment preferentially increased the trans-associated 9- OOH(EE) species, together with 13-OOH products (**Figure 5D**). These findings support the conclusion that trans fatty acids are incorporated into membrane phospholipids and undergo radical-mediated oxidation during ferroptosis, generating oxidation products with a stereochemical signature consistent with the precursor fatty acid.

Petroselaidic acid produced comparatively little accumulation of the quantified linoleate-derived phosphatidylcholine hydroperoxides (**Figure 5D**). However, interpretation of these results is limited by the targeted nature of the assay, which quantified a defined subset of oxidized phosphatidylcholine species for which authentic standards were available. Given our earlier finding that petroselaidic acid undergoes SCD-dependent conversion and promotes extensive phospholipid remodeling, it remains possible that oxidation of additional phospholipid classes or molecular species, including remodeled trans-containing phospholipids not captured in the present analysis, contributes to its ferroptosis-sensitizing activity.

We also examined oxidation of arachidonate-containing phospholipids. Although several oxidized arachidonate-containing phosphatidylcholine species increased following fatty acid treatment, the overall pattern of arachidonate hydroperoxides was broadly similar between linoleic acid- and linoelaidic acid-treated cells (**Figure 5E**). Thus, the principal biochemical distinction between cis and trans fatty acids was not a global increase in phospholipid oxidation but rather the generation of unique oxidation products derived directly from trans-containing phospholipids.

Together, our findings demonstrate that incorporation of trans fatty acids into membrane phospholipids gives rise to a chemically distinct repertoire of oxidized phospholipid species during ferroptosis. While linoelaidic acid directly generates abundant trans-specific phospholipid hydroperoxides, the comparative lack of oxidation detected following petroselaidic acid treatment suggests that its ferroptosis-sensitizing activity likely reflects broader metabolic remodeling of the membrane lipidome that is not reflected in our targeted analysis. Further characterization of oxidized phospholipid species beyond the targeted phosphatidylcholine hydroperoxides analyzed here will be important to fully define the downstream oxidation products generated following petroselaidic acid metabolism.

### Trans fatty acids promote the accumulation of ferroptosis-sensitizing phospholipid species

To define how trans fatty acid metabolism remodels the phospholipid landscape to generate a membrane composition that favors ferroptosis, H460 cells were treated with linoleic acid, linoelaidic acid, or petroselaidic acid for 24 hours prior to GC-FID fatty acid profiling and phospholipid LC-MS/MS lipidomic analyses. Fatty acid profiling demonstrated that each treatment remodeled the cellular fatty acid pool in distinct ways (**Figure 6A**). Consistent with direct incorporation, both linoleic acid and linoelaidic acid markedly increased the overall abundance of PUFAs, whereas petroselaidic acid predominantly increased the MUFA pool and resulted in a smaller increase in PUFAs (**Figure 6A**). Despite these differences in overall fatty acid composition, both trans fatty acids robustly sensitized cells to ferroptosis more strongly than cis PUFA linoleic acid (**Figure 6B**). These results indicate that changes in bulk PUFA content alone cannot predict susceptibility to ferroptosis; the particulars of fatty acid metabolism and incorporation into cellular lipids dictate the oxidizability of cellular membranes.

**Figure 6.**
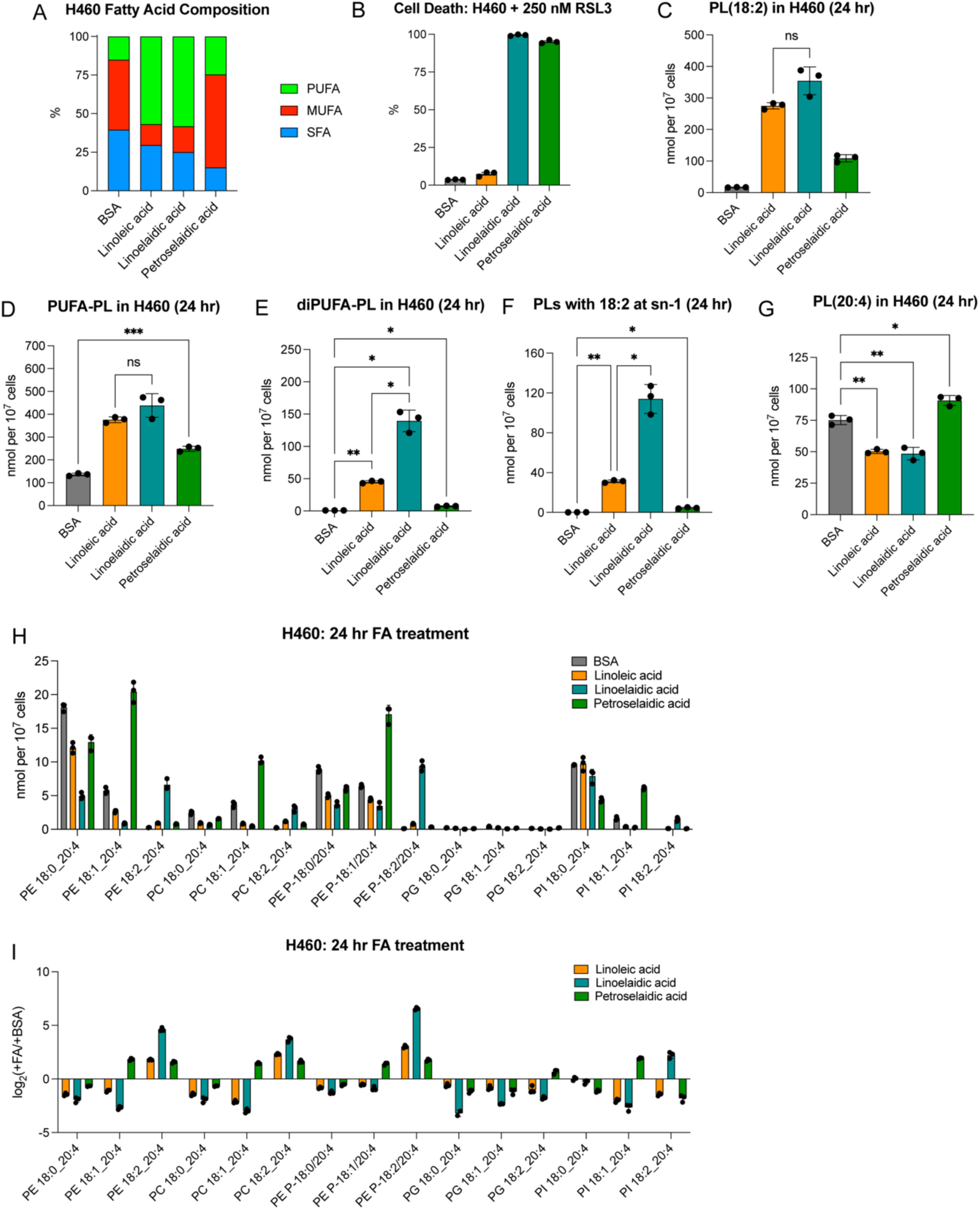
Trans fatty acids remodel membrane phospholipids to generate distinct ferroptosis-susceptible lipid species. (**A**) Overall lipid composition of H460 cells treated with BSA or 100 µM indicated fatty acid for 24 hours as determined by GC-FID. (**B**) Cell death of H460 cells treated as in (**A**) then treated with 250 nM RSL3 for 24 hours. (**C**) Quantification of 18:2-containing phospholipids by shotgun lipidomics for H460 cells treated as in (**A**). (**D**) Quantification of PUFA-containing phospholipids by shotgun lipidomics for H460 cells treated as in (**A**). (**E**) Quantification of diPUFA phospholipids by shotgun lipidomics for H460 cells treated as in (**A**). (**F**) Quantification of phospholipids containing 18:2 at the sn-1 position by shotgun lipidomics for H460 cells treated as in (**A**). (**G**) Quantification of 20:4-containing phospholipids by shotgun lipidomics for H460 cells treated as in (**A**). (**H**) Quantification of phospholipids containing 20:4 and an 18-carbon fatty acyl tail by shotgun lipidomics for H460 cells treated as in (**A**). (**I**) Enrichment and dis-enrichment calculated by log2(fatty acid treatment/BSA treatment) of phospholipids in (**H**) for H460 cells treated as in (**A**).

We next examined incorporation into membrane phospholipids using lipidomics. Total 18:2- containing phospholipids increased similarly following linoleic acid and linoelaidic acid treatment, whereas petroselaidic acid produced substantially lower amounts of 18:2 phospholipids (**Figure 6C**). Similarly, total PUFA-containing phospholipids were comparably elevated following linoleic acid and linoelaidic acid treatment (**Figure 6D**). These observations suggest that the enhanced ferroptosis sensitization induced by linoelaidic acid is not simply explained by greater overall incorporation into PUFA-containing phospholipids. Instead, analysis of individual phospholipid subclasses revealed marked differences in phospholipid remodeling. Linoelaidic acid treatment significantly increased the abundance of diPUFA-containing phospholipids (**Figure 6E**), lipid species that have recently emerged as particularly susceptible substrates for ferroptotic lipid peroxidation^15^. In addition, linoelaidic acid preferentially increased phospholipids containing an 18:2 fatty acid at the *sn*-1 position (**Figure 6F**), an unusual phospholipid architecture that is uncommon under basal conditions and consistent with the incorporation of trans 18:2 PUFA into the *sn*-1 position of membrane phospholipids.

Petroselaidic acid produced a different remodeled lipidome pattern. Rather than increasing diPUFA phospholipids, petroselaidic acid increased arachidonate-containing phospholipids (**Figure 6G**) and selectively remodeled multiple phospholipid classes, including phosphatidylethanolamine, phosphatidylcholine, phosphatidylinositol, and phosphatidylglycerol species (**Figure 6H,I**). Notably, several of the most strongly increased phospholipids were consistent with species containing an 18:1 fatty acid paired with arachidonic acid (e.g., PE 18:1_20:4), raising the possibility that petroselaidic acid is incorporated at the *sn*-1 position while arachidonic acid occupies the *sn*-2 position (**Figure 6H,I**). Such phospholipid architectures provide abundant polyunsaturated substrates for ferroptotic lipid peroxidation while simultaneously incorporating the trans fatty acid into membrane phospholipids.

Overall, these data demonstrate that structurally distinct trans fatty acids converge on ferroptosis sensitization through different phospholipid remodeling pathways. Whereas linoelaidic acid preferentially promotes the accumulation of diPUFA phospholipids and trans 18:2-containing phospholipids, petroselaidic acid remodels membranes toward arachidonate-rich phospholipid species. Despite these distinct metabolic routes, both trans fatty acids expand the pool of phospholipid species predicted to be highly susceptible to ferroptotic lipid peroxidation.

## DISCUSSION

Ferroptosis is fundamentally governed by the interplay between lipid peroxidation chemistry and membrane lipid composition. Previous studies have established fatty acid chain length, degree of unsaturation, phospholipid class, and phospholipid remodeling pathways as major determinants of ferroptosis susceptibility by regulating the abundance of oxidation-prone membrane phospholipids^4,10^. Here, we identify fatty acid double-bond geometry as an additional structural determinant of ferroptosis sensitivity. Through an unbiased functional fatty acid screen, we unexpectedly identified trans fatty acids as potent ferroptosis sensitizers and demonstrate that cis-trans double bond geometry determines fatty acid metabolic fate, remodels membrane phospholipid composition, and promotes the formation of ferroptosis-susceptible membrane lipid species. A central finding of this study is that fatty acid stereochemistry influences ferroptosis primarily through its effects on lipid metabolism rather than simply through intrinsic lipid chemistry. Previous work has largely focused on the inherent susceptibility of PUFAs to radical-mediated oxidation, whereas our findings demonstrate that the metabolic processing of a fatty acid is equally important in determining whether it ultimately contributes to membrane lipid peroxidation. Thus, in addition to chain length and degree of unsaturation, double-bond geometry represents a previously unappreciated structural feature that shapes membrane composition and ferroptosis sensitivity (**Figure 7**).

**Figure 7.**
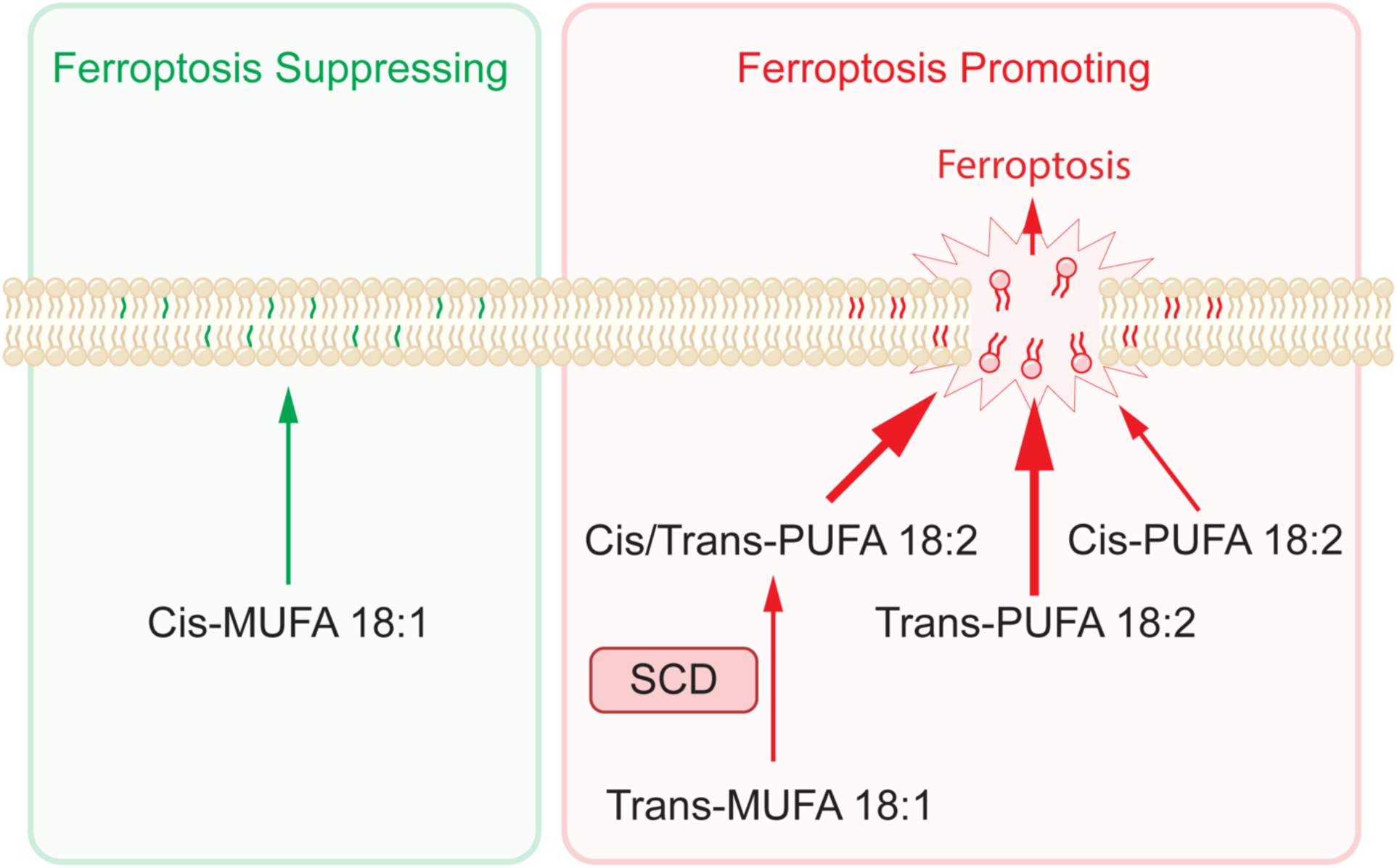
Model: Double-bond geometry programs fatty acid metabolic fate to regulate membrane composition and ferroptosis sensitivity. Cis 18:1 MUFAs such as oleic acid and petroselinic acid contain central double bonds that result in a singly-kinked structure that suppresses ferroptosis. Petroselaidic acid, a trans 18:1 MUFA with a structure permissible to Δ9-desaturation by SCD, undergoes conversion to a mixed trans/cis 18:2 PUFA and promotes ferroptosis. The trans 18:2 PUFA linoelaidic acid and its mixed cis/trans 18:2 PUFA stereoisomers unexpectedly enhance ferroptosis to a greater degree than the essential omega-6 cis 18:2 PUFA linoleic acid through increased incorporation at the sn-1 position of phospholipids, resulting in an increase in the formation of diPUFA phospholipids.

Our findings illustrate this principle through two mechanistically distinct trans fatty acids. Linoelaidic acid was efficiently incorporated into phospholipids and promoted the accumulation of diPUFA phospholipids, phospholipids enriched in 18:2 at the *sn*-1 position, and trans-specific phospholipid hydroperoxides following GPX4 inhibition. In contrast, the trans MUFA petroselaidic acid underwent SCD-dependent desaturation to generate a trans PUFA and promoted remodeling toward arachidonate-containing phospholipid species. The identification of the trans MUFA petroselaidic acid as a potent ferroptosis sensitizer was an unexpected and striking finding since cis MUFAs are established suppressors of ferroptosis. Our findings also distinguish non-conjugated trans fatty acids from previously described conjugated PUFAs, which sensitize to ferroptosis because their conjugated double-bond systems are intrinsically highly susceptible to radical-mediated oxidation^48–50^. In contrast, biochemical studies of lipid oxidation have shown that trans 18:2 PUFAs exhibit slower oxidation kinetics than those of their cis counterparts^47^ and that 20:4 PUFAs that contain a trans bond suppress ferroptosis^51^. Consistent with these observations, our data support a model in which trans fatty acids do not promote ferroptosis because they are inherently more oxidizable. Rather, trans fatty acids are metabolized differently, leading to phospholipid remodeling that expands the abundance of ferroptosis-susceptible membrane lipids available for peroxidation.

Several observations suggest that additional mechanisms remain to be defined, and our study has certain limitations. Although linoelaidic acid generated abundant trans-specific phospholipid hydroperoxides, oxidation products derived from petroselaidic acid were comparatively modest in our targeted phospholipid analyses despite its robust ferroptosis-sensitizing activity. One explanation is that the targeted lipidomics performed here captured only a subset of oxidized phosphatidylcholine species for which authentic standards were available. Our phospholipid remodeling data, together with the detection of additional putative oxidized phospholipid species, raise the possibility that oxidation of remodeled phosphatidylethanolamine or other phospholipid classes contributes to the activity of petroselaidic acid. Likewise, although our lipidomics data are consistent with preferential enrichment of trans fatty acids at the *sn*-1 position of phospholipids, definitive structural assignment of individual phospholipid molecular species will require additional analytical approaches and authentic standards.

Beyond ferroptosis, our findings have broader implications for lipid biology. Trans fatty acids arise from both dietary intake and endogenous lipid isomerization during oxidative stress^52^, suggesting that fatty acid stereochemistry may represent a previously underappreciated mechanism for regulating membrane composition and function. Our findings may also have therapeutic implications. SCD expression is elevated in many cancers and has emerged as a determinant of ferroptosis sensitivity through its role in fatty acid metabolism^14,53^. The requirement for SCD-dependent metabolic activation of petroselaidic acid raises the intriguing possibility that tumors with high SCD activity may be selectively susceptible to trans PUFA-mediated ferroptosis sensitization. More broadly, our work expands the structural principles governing membrane lipid homeostasis by demonstrating that stereochemical differences, independent of chain length or degree of unsaturation, can profoundly alter fatty acid metabolism, phospholipid remodeling, and downstream cellular physiology.

In summary, we propose a model in which cis-trans double-bond geometry determines the metabolic fate of fatty acids, leading to distinct phospholipid remodeling pathways that converge on the formation of ferroptosis-susceptible membrane lipids. Rather than altering the intrinsic chemical susceptibility of fatty acids to oxidation, double-bond geometry directs the generation of membrane phospholipid species that are poised to undergo ferroptotic lipid peroxidation. Our work suggests that the ferroptosis sensitivity of a fatty acid cannot be predicted solely from its chemical structure but instead emerges from the interplay between fatty acid stereochemistry, cellular metabolism, and membrane phospholipid remodeling. Together, these findings establish fatty acid stereochemistry as a previously unrecognized determinant of membrane composition and ferroptotic cell fate and suggest that the biological activities of fatty acids emerge not simply from their chemical structures but from the metabolic pathways through which those structures are interpreted by the cell.

## ACKNOWLEDGEMENTS

This research was supported by grants from the National Institutes of Health to J.A.O. (R01CA305423 and R01CA276207) and A.G. (R01DK095045 and R01DK099465). S.K. and Y.O. were supported by the JSPS KAKENHI Grant Number 25K09014. C.A.H. was supported in part by a University of California Cancer Research Coordinating Committee (CRCC) predoctoral fellowship award and in part by a predoctoral fellowship from the Northern California Chapter of the Achievement Rewards for Career Scientists (ARCS) Foundation. We thank Kartoosh Heydari, Melaine Delcroix, and Harman Dhaliwal of the UC Berkeley Cancer Research Laboratory (CRL) Flow Cytometry Core Facility for training on and use of flow cytometry analyzers. We thank Alison Killilea and Sara Sosa of the UC Berkeley Cell Culture Facility which is supported by the University of California Berkeley for providing cell lines used in this study (RRID: SCR_017924). We thank Alice Refermat and Pingping He of the High-Throughput Screening Facility (HTSF) at UC Berkeley. This work was performed in part in the HTSF, which provided the Perkin Elmer Opera Phenix. Research reported in this publication was supported by the Office of the Director, National Institutes of Health, under Award Number S10OD021828. The content is solely the responsibility of the authors and does not necessarily represent the official views of the National Institutes of Health.

## AUTHOR CONTRIBUTIONS

C.A.H. and J.A.O. conceived of the project, designed the experiments, and wrote the majority of the manuscript. C.A.H. performed the majority of experiments and analyses. S.K. and Y.O. performed lipid hydroperoxide quantification. E.M.P. performed the lipid droplet imaging and quantification. A.G., J.L.P. and D.B. provided fatty acid reagents for the FALCON screen. All authors read, edited, and contributed to the manuscript.

## METHODS

### Cell lines and culture conditions

All cells were maintained at 37°C, 5% CO_2_ in a humidified incubator. With the exception of Northern elephant seal primary endothelial cells which were a gift from the José Pablo Vázquez-Medina lab, all cell lines were obtained from the Cell Culture Facility (CCF) at UC Berkeley. H441, H460, H520, H1703, and H2291 non-small cell lung cancer lines were grown in RPMI 1640 with 10% FBS. Seal primary endothelial cells were grown in low-glucose DMEM supplemented with 10% FBS, 1% NEAA, 1% HEPES, and 1% antibiotic/antimycotic. All other cell lines were grown in DMEM with 10% FBS. All cell lines were confirmed to be mycoplasma free by regular PCR tests.

### Preparation of BSA-conjugated fatty acid stocks

Linoleic acid (Cayman Chemical #90150), linoelaidic acid (Cayman Chemical #90160), petroselaidic acid (Cayman Chemical #20026), petroselinic acid (Cayman Chemical #20024), arachidonic acid (Cayman Chemical #90010), 9(Z),12(E)-octadecadienoic acid (Larodan #10- 1852-2), and 9(E),12(Z)-octadecadienoic acid (Cayman Chemical #10005146) were diluted in 100% molecular-biology-grade ethanol to a stock concentration of 200 mM. Fatty acids in ethanol were then further diluted to 10 mM with 10% fatty acid-free BSA (Fisher Scientific #BP9704) in borosilicate vials and incubated at 37°C overnight. BSA-conjugated fatty acid stocks were aliquoted and stored in borosilicate vials at −80°C.

### Dose-response and lethal fraction assays

Cell lines were lentivirally transduced to express mCherry-NLS for lethal fraction calculations in dose-response assays. Cells were seeded into flat black clear-bottom 96-well plates (Corning #3904). Drug/fatty acid treatments were added at 2x concentration in an equal volume of media containing Sytox Green Dead Cell Stain (Thermo Fisher #S34860) at 2x concentration (30 nM final concentration). Drugs and inhibitors used in these assays were purchased from Cayman Chemical: (1S,3R)-RSL3 (#19288), Z-VAD-FMK (#14463), Necrostatin-1 (#11658), Ferrostatin-1 (#17729), idebenone (#15475), deferoxamine mesylate (DFO, #14595), ML-162 (#20455), ML- 210 (#23282), Erastin2 (#27087), Imidazole Ketone Erastin (IKE, #27088), L-Buthionine-(S,R)- Sulfoximine (BSO, #14484), FINO_2_ (#25096), FIN56 (#25180), FSEN1 (#38025), CAY10566 (#10012562), SC-26196 (#10792), A-922500 (#10012708), and PF-06424439 (#17680). Cell death was monitored over time by the Incucyte S3 live-cell analysis system (Sartorius). For fatty acid pre-treatments, 2x fatty acid in media was first added to plates in 50 µL per well; 50 µL cells in suspension were then seeded into fatty acid-containing wells. Lethal fraction was calculated by dividing the number of Sytox Green positive objects by the sum of red objects (mCherry-positive nuclei) and Sytox Green Positive objects, minus the number of overlapping green and red objects.

### FALCON screen for fatty acid synergy with RSL3

FALCON library plates were prepared as previously described. Briefly, 10 mM stocks of each fatty acid in DMSO were dried in vacuo and conjugated to fatty acid-free BSA. BSA-conjugated fatty acid was distributed into deep well 384-well plates and once again dried. One day prior to treatment, H460 and U-2 OS cells were seeded into 384-well plates at 1200 or 1500 cells per well, respectively. At the time of treatment, a 1:1 mixture of RPMI 1640 + 10% FBS and DMEM + 10% FBS containing 60 nM Sytox Green and either DMSO or 2 µM RSL3 was added to the dried BSA-conjugated fatty acid plates to a final concentration of 1 mM FFA. Plates were incubated at 37°C for 1 hour and shaken on a plate shaker at room temperature for 30 min. Dissolved fatty acids were serially diluted in DMSO- or 2 µM RSL3-containing Sytox Green media and added to cell plates at 2x concentration. Cell death was monitored via Incucyte S3 live-cell analysis system (Sartorius).

### GC–FID for absolute fatty acid quantification

H460 cells were treated with BSA or 100 µM fatty acid for 24 hours, collected via scraping into cold DPBS and pelleted at 500xg for 5 min at 4°C. The fatty acid compositions of the pelleted cells were analyzed at OmegaQuant (Sioux Falls, SD) via GC–FID. Total cell fatty acids were hydrolyzed and methylated into fatty acid methyl esters (FAMEs) using an internal protocol. FAMEs were separated on a GC2010 gas chromatograph set-up (Shimadzu) equipped with an SP2560 fused-silica capillary column (100 m, 0.25-mm internal diameter, 0.2-µm film thickness; Supelco). The fatty acids were identified based on the characteristic elution times of a fatty acid standard mixture (GLC OQ-A, Nu-Check Prep), quantified based on external calibration, and normalized to cell count.

### Immunoblots

Cells rinsed with DPBS, collected by scraping, and pelleted at 500xg for 5 min at 4°C. Cell pellets were lysed in RIPA buffer with EDTA-free Pierce Protease and Phosphatase Inhibitor mini tablets (Thermo Scientific #A32961) and sonicated at 15% amplitude for 30 s. Protein concentrations were determined using the bicinchoninic acid (BCA) protein assay (Thermo Scientific #23227). Even amounts of protein by mass were combined with 4x Laemmli buffer (Bio-Rad Laboratories #1610747) supplemented with beta-mercaptoethanol to a final concentration (v/v) of 5%. Samples were loaded on 4–20% polyacrylamide gradient gels (Bio-Rad Laboratories #4561094) and transferred onto nitrocellulose membranes (Bio-Rad Laboratories #1620115). Membranes were blocked in EveryBlot Blocking Buffer (Bio-Rad Laboratories #12010020) at room temperature on a platform rocker for 10 min then rinsed with PBS containing 0.1% Tween 20 (PBS-T). Membranes were incubated overnight at 4 °C in PBS-T containing 3% BSA and primary antibodies (1:1000 dilution of mouse anti-SCD1 [CD.E10], Abcam #ab19862; 1:5000 dilution of rabbit anti-beta-actin (13E5), Cell Signaling Technology #4970). After three 10 minute rinses with room temperature PBS-T on a platform rocker, membranes were transferred to PBS-T containing 3% BSA and a 1:10,000 dilution of fluorescent secondary antibodies (IRDye 800CW Donkey anti-Mouse LI-COR #926-32212 or IRDye 800CW Donkey anti-Rabbit LI-COR #926-68073) and rocked at room temperature for 1 hr. After three 10 minute rinses with PBS-T, immunoblots were imaged on a LI-COR Odyssey imager (LI-COR Biosciences).

### Lipid droplet imaging and quantification

Lipid droplet imaging was performed using a high-throughput Opera Phenix confocal microscope (Revvity), followed by quantification with the Harmony High Content Image Analysis software (Revvity) using a user-defined pipeline. H460 cells were seeded in flat black clear-bottom 96-well plates (Corning, 3904) at a density of 3,500 cells/well. After 24 hours, cells were treated with 100 µM linoleic acid, linoelaidic acid, or petroselaidic acid complexed to fatty acid-free BSA, either alone or in combination with 45 µM A- 922500 DGAT1 inhibitor (Cayman Chemical #10012708) and 30 µM PF-06424439 DGAT2 inhibitor (Cayman Chemical #17680). A BSA-only, vehicle control was also included. Following 24 hours of treatment, cells were fixed with 4% paraformaldehyde, washed twice with PBS, and incubated with 1 µg/mL DAPI and 1 µM BODIPY 493/503 (Thermo Fisher Scientific, D3922) for 30 min at room temperature in the dark. Cells were then washed twice with PBS and imaged using a 40x water-immersion objective. Each biological replicate consisted of 3 technical replicates (3 wells per condition), with 15 fields of view acquired per well.

Lipid droplets were quantified from confocal images using maximum intensity projections of z-stacks for each field of view. The DAPI (blue) channel was first used to segment nuclei via the “Find Nuclei” function, and only nuclei not touching the edges of the field of view were selected for downstream analysis. The BODIPY 493/503 (green) channel was then used to segment the cytoplasm via “Find Cytoplasm,” excluding the nuclear region and defining a cytoplasmic mask immediately surrounding the nucleus. The same channel was subsequently used to identify puncta corresponding to lipid droplets using the “Find Spots” function, with further optimization for punctal intensity and background correction. The mean number of lipid droplets per cytoplasmic area was quantified for each condition and plotted using Prism (GraphPad). Statistical analysis was performed using a one-way ANOVA with multiple comparisons (Dunnett’s multiple comparisons test).

### BODIPY 493/503 staining for flow cytometry

Sub-confluent H460 cells in 6-well plates were treated with 100 µM fatty acid or an equivalent amount of fatty acid-free BSA in complete media with or without 45 µM A-922500 DGAT1 inhibitor and 30 µM PF-06424439 DGAT2 inhibitor for 24 hours. Cells were then rinsed with DPBS and stained with 1 µM BODIPY 493/503 in Opti-MEM (Gibco, 31985070) at 37°C in a dark tissue-culture incubator for 30 min. Following staining, cells were collected for analysis by flow cytometry.

### BODIPY C11 staining for flow cytometry

Sub-confluent H460 cells in 6-well plates were pre-treated with 100 µM fatty acid or an equivalent amount of fatty acid-free BSA in complete media with or without 5 µM ferrostatin-1 for 20 hours. After pre-treatment, cells were rinsed with DPBS, and the media was replaced with Opti-MEM containing DMSO, 100 nM RSL3, or 100 nM RSL3 and 5 µM ferrostatin-1. After 2 hours of treatment, C11 BODIPY 581/591 (Invitrogen, D3861) suspended in OptiMEM was added directly to the media in each well at 2x concentration for a final concentration of 1 µM and allowed to stain at 37°C in a dark tissue-culture incubator for 30 min. Following staining, cells were collected for analysis by flow cytometry.

### Liperfluo staining for flow cytometry

Sub-confluent H460 cells in 6-well plates were pre-treated with 100 µM fatty acid or an equivalent amount of fatty acid-free BSA in complete media with or without 5 µM ferrostatin-1 for 20 hours. After pre-treatment, cells were rinsed with DPBS, and the media was replaced with serum-free OptiMEM containing 1 µM Liperfluo (Dojindo #L248) and either DMSO, 5 µM ferrostatin-1, 100 nM RSL3, or 100 nM RSL3 and 5 µM ferrostatin-1 for 2 hours. Following staining, cells were collected for analysis by flow cytometry.

### Flow cytometry

Cells were washed twice in DPBS and dissociated using TrypLE Express (Gibco, 12605010). Cells were pelleted at 500xg, 4°C for 2 min, the supernatant was removed, and cells were rinsed with cold PBS. The cells were pelleted once again at 500xg, 4°C for 2 min before resuspension in 300 µL cold PBS for analysis.

For all flow cytometry assays, fluorescence was analyzed on an LSR Fortessa analyzer (BD Biosciences) using BD FACSDiva v.6.2 (BD Biosciences). The following filter sets were used: FITC (BODIPY 493/503, BODIPY C11 oxidized channel, Liperfluo) and PE-Texas-Red (BODIPY C11 non-oxidized channel). FlowJo v.10 software (BD Biosciences) was used to quantify fluorescence and generate representative histograms.

### LC-MS/MS detection of phospholipid hydroperoxides

60 million H460 cells were plated into media containing BSA or 100 µM linoleic acid, linoelaidic acid, or petroselaidic acid in 500 cm² square TC-treated culture dishes (Corning #431110). After 20 hours (>1 doubling time), cells were rinsed twice with DPBS and scraped into cold DPBS then pelleted at 500xg, 4°C for 5 min. After removal of the supernatant, pellets were resuspended in 2 mL DPBS and transferred to solvent-compatible 2.0 mL Safe-Lock tubes (Eppendorf #0223363352) and stored at −80°C. Cell pellets were subjected to lipid extraction using the Folch method, followed by solid-phase extraction to isolate the phospholipid fraction, as described previously^54^. The purified phospholipid fraction was dissolved in 200 μL of methanol, and a 10 μL aliquot was subjected to LC–MS/MS analysis as described below.

LC–MS/MS analysis was performed using an ExionLC system coupled to a QTRAP 6500 mass spectrometer (SCIEX, Tokyo, Japan). Chromatographic separation was achieved on a COSMOSIL 5C18-MS-II column (2.0 × 150 mm; Nacalai Tesque, Inc., Kyoto, Japan) maintained at 40°C. The mobile phases consisted of water (A) and methanol (B). The flow rate was 0.30 mL/min. The initial mobile phase composition was 85% B. The gradient program was as follows: 85% B initially, increased to 92.5% B at 15.0 min, to 100% B at 15.1 min, held at 100% B until 25.0 min, and returned to 85% B at 25.1 min for column re-equilibration. A post-column solution of 2 mM sodium acetate in methanol was introduced at a flow rate of 0.01 mL/min to promote the formation of sodium adduct ions, thereby enabling all precursor ions to be detected as [M+Na]⁺ species. Multiple reaction monitoring (MRM) transitions and optimized MS parameters are listed in **Supplementary Table 1**. Quantification was performed using the external calibration method. Authentic standards of PC(16:0/18:2-OOH) were prepared as previously reported^55^, and the same synthetic strategy was applied to prepare PC(16:0/20:4-OOH) standards.

### Targeted lipidomic quantification of phospholipid species

H460 cells were treated with BSA or 100 µM fatty acid for 24 hours then collected via scraping into cold DPBS and pelleted at 500xg for 5 min. Lipidomic analysis was performed at the UCLA Lipidomics Core. Cell pellets were homogenized in PBS using 3 10-second rounds of ceramic bead-based homogenization. Lipids were extracted from the homogenates using a modified Bligh and Dyer method and combined with 70 internal lipid standards (AB Sciex, Avanti). Following two rounds of organic solvent extraction, the combined solvent layers were dried in vacuo (35°C, 90 min in total). The dried lipids were resuspended in methanol/dichloromethane with ammonium acetate and transferred to analysis-grade vials. Lipid species were quantified using a Sciex 5500 instrument equipped with a differential mobility device (Lipidyzer), calibrated with EquiSPLASH standards, detecting up to 1450 lipid species across 17 subclasses using a predefined method. Data were processed via an in-house platform and normalized to cell count to ensure accuracy.

### Statistical analysis

Statistical analyses were performed with GraphPad Prism 11 using ordinary one-way analysis of variance (ANOVA) and Dunnett’s multiple comparisons test or Brown-Forsythe and Welch ANOVA tests with Dunnett’s T3 multiple comparisons test with individual variances computed for each comparison (***P < 0.001, **P < 0.01, *P < 0.05). Error bars in dose-response plots represent standard error of the mean (SEM); error bars in bar graphs represent standard deviation.

---

**Supplementary Figure S1.**
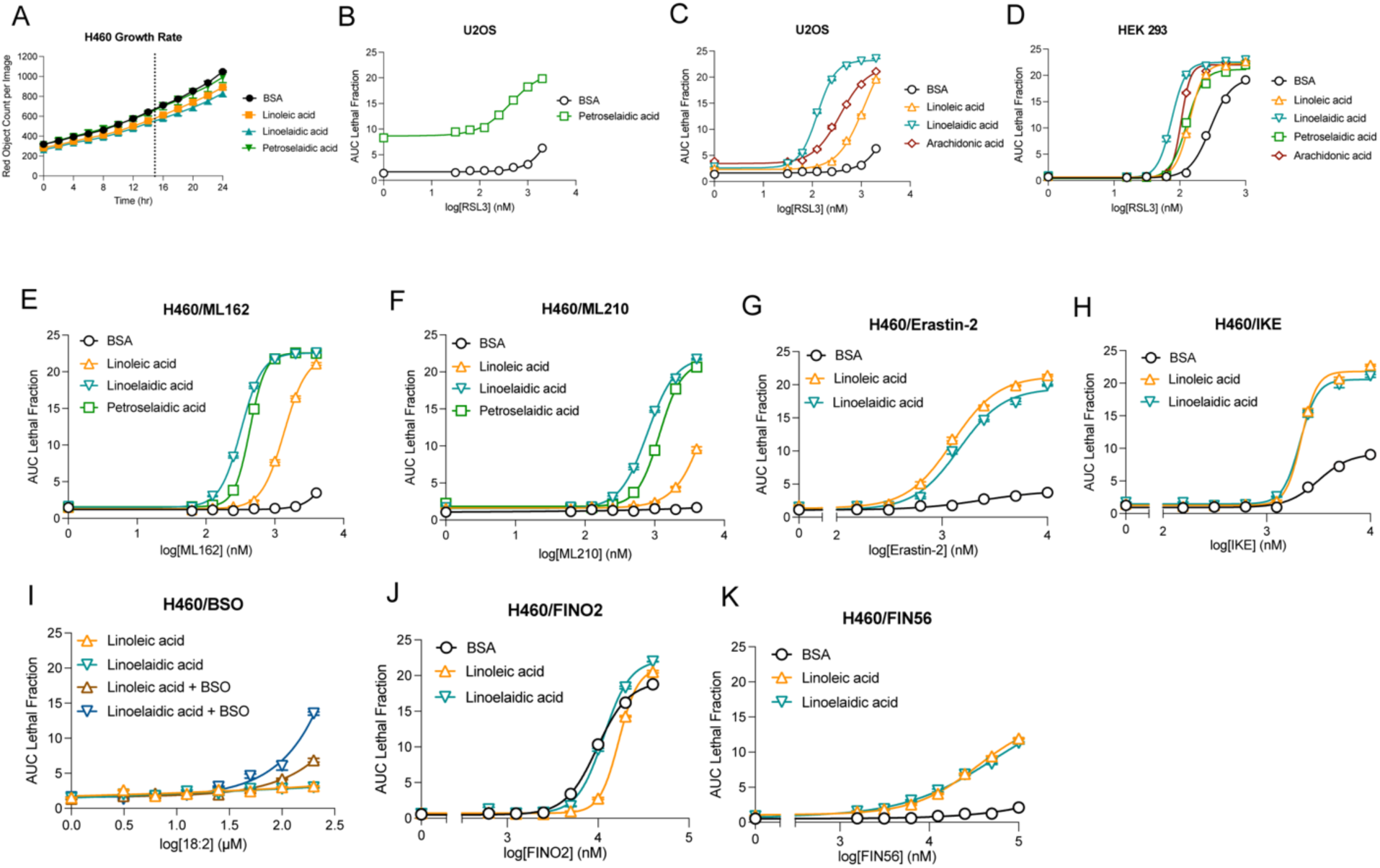
Trans fatty acids promote sensitivity to ferroptosis-inducing drugs. (**A**) H460 nucleus count per image over time with 100 µM indicated fatty acid treatments. Vertical dashed line represents population doubling (approximately 15 hours). (**B**) RSL3 dose response of U-2 OS cells pre-treated with either BSA or 100 µM petroselaidic acid for 24 hours. (**C**) RSL3 dose response of U-2 OS cells pre-treated with either BSA or 100 µM indicated fatty acids for 24 hours. (**D**) RSL3 dose response of HEK 293 cells pre-treated with either BSA or 100 µM indicated fatty acids for 24 hours. (**E**) ML162 dose response of H460 cells pre-treated with either BSA or 100 µM indicated fatty acids for 24 hours. (**F**) ML210 dose response of H460 cells pre-treated with either BSA or 100 µM indicated fatty acids for 24 hours. (**G**) Erastin-2 dose response of H460 cells pre-treated with either BSA or 200 µM indicated fatty acids for 24 hours. (**H**) IKE dose response of H460 cells pre-treated with either BSA or 200 µM indicated fatty acids for 24 hours. (**I**) Fatty acid dose response of H460 cells pre-treated with indicated fatty acids for 24 hours followed by treatment with 500 µM BSO. (**J**) FINO2 dose response of H460 cells pre-treated with either BSA or 200 µM indicated fatty acids for 24 hours. (**K**) FIN56 dose response of H460 cells pre-treated with either BSA or 200 µM indicated fatty acids for 24 hours.

**Supplementary Figure S2.**
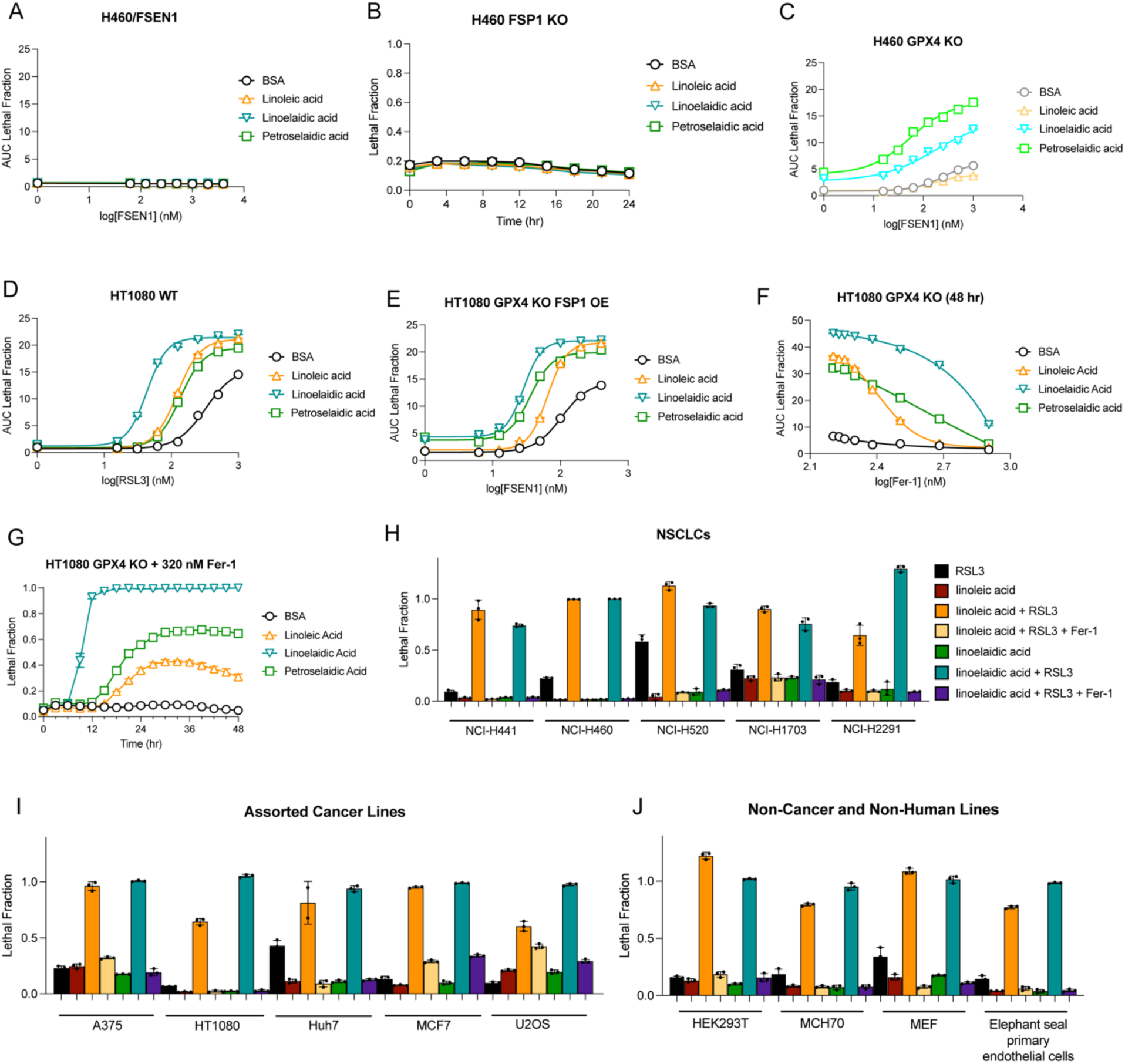
Trans fatty acids sensitize multiple cell lines to GPX4 inhibition. (**A**) FSEN1 dose response of H460 cells pre-treated with either BSA or 100 µM indicated fatty acid for 24 hours. (**B**) Lethal fraction of H460 FSP1 KO cells over time after 24 hour BSA or 100 µM fatty acid treatment. (**C**) FSEN1 dose response of H460 GPX4 KO cells pre-treated with either BSA or 100 µM indicated fatty acid for 24 hours. (**D**) RSL3 dose response of HT1080 cells pre-treated with either BSA or 100 µM indicated fatty acids for 24 hours. (**E**) FSEN1 dose response of HT1080 GPX4 KO FSP1 OE cells pre-treated with either BSA or 100 µM indicated fatty acids for 24 hours. (**F**) 48 hour Ferrostatin-1 washout dose response of HT1080 GPX4 KO cells pre-treated with either BSA or 100 µM indicated fatty acids for 24 hours. (**G**) Lethal fraction over time after ferrostatin-1 dilution to <320 nM in HT1080 GPX4 KO cells pre-treated with BSA or 100 µM indicated fatty acids. (**H**) Lethal fraction of non-small cell lung cancer cell lines (NSCLCs) with indicated treatments. Linoleic acid and linoelaidic acid pre-treatments at 200 µM for 24 hours. Ferrostatin-1 at 5 µM. RSL3 doses: H441, 50 nM; H460, 1 µM; H520, 1 µM; H1703, 25 nM; H2291, 100 nM. (**I**) Lethal fraction of assorted cancer cell lines with indicated treatments. Linoleic acid and linoelaidic acid pre-treatments at 200 µM for 24 hours. Ferrostatin-1 at 5 µM. RSL3 doses: A375, 100 nM; HT1080, 100 nM; Huh7, 250 nM; MCF7, 500 nM; U2OS, 250 nM. (**J**) Lethal fraction of non-cancer and non-human cell lines with indicated treatments. Linoleic acid and linoelaidic acid pre-treatments at 200 µM for 24 hours. Ferrostatin-1 at 5 µM. RSL3 doses: HEK293T, 50 nM; MCH70, 100 nM; MEF, 100 nM; elephant seal primary endothelial cells, 500 nM.

**Supplementary Figure S3.**
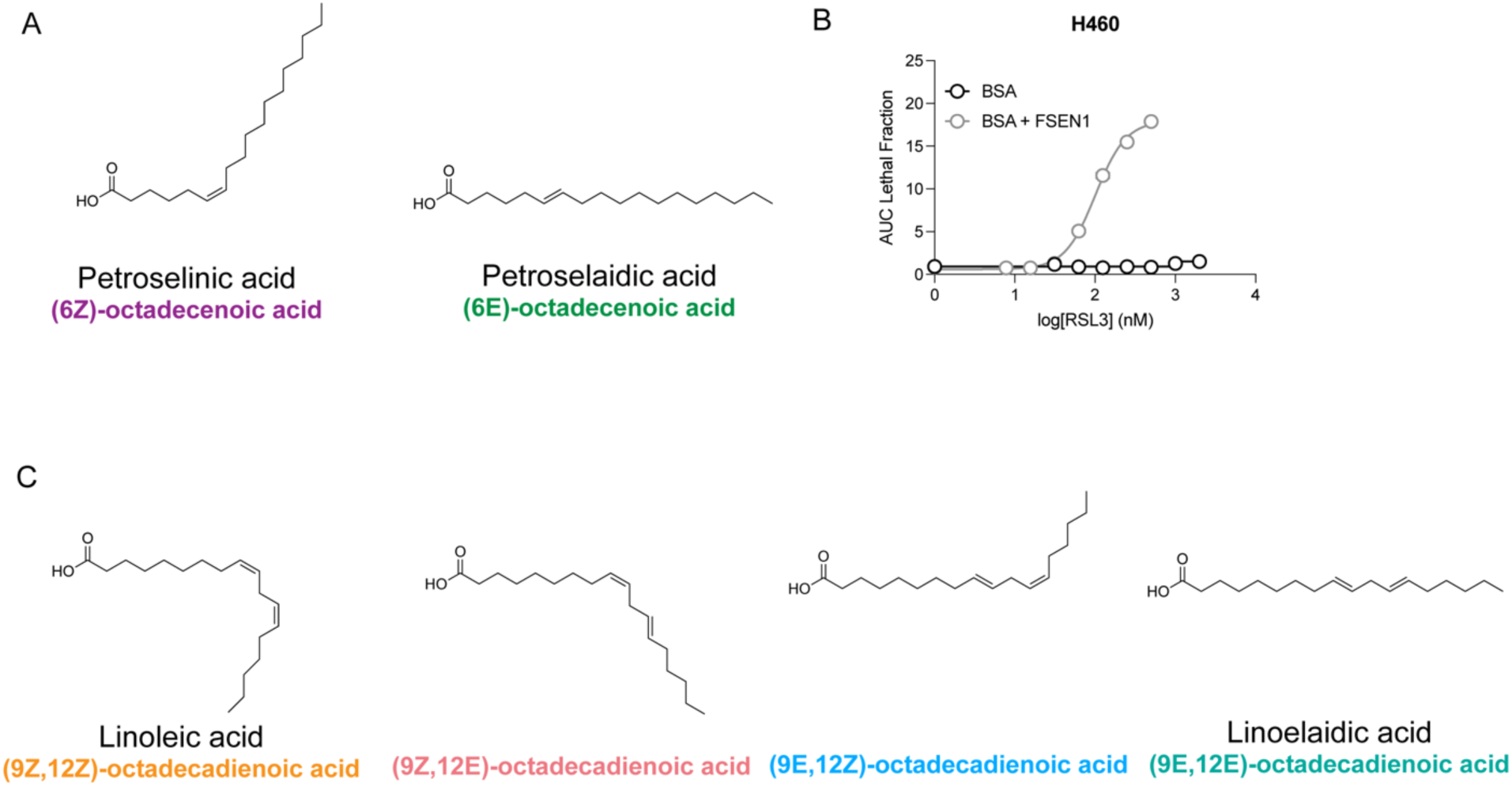
**Fatty acid stereoisomers have distinct structures.**(**A**) Structures of cis MUFA petroselinic acid compared to trans MUFA petroselaidic acid. (**B**) RSL3 dose response in the presence or absence of 1 µM FSEN1 of H460 cells pre-treated with BSA for 24 hours. (**C**) Structures of cis PUFA linoleic acid and trans PUFA linoelaidic acid compared to their mixed cis,trans and trans,cis stereoisomers.

**Supplementary Figure S4.**
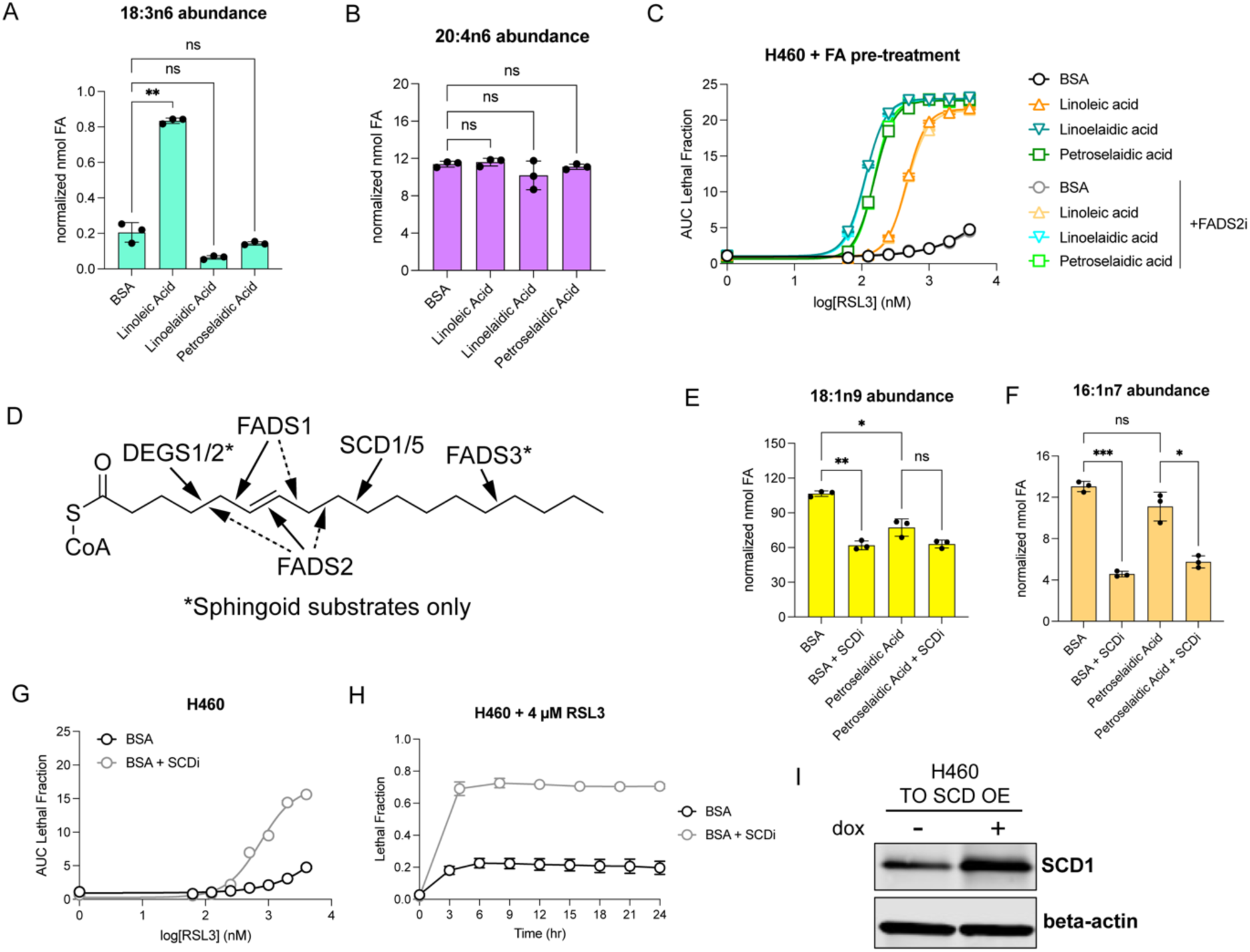
Fatty acids are regio-specifically desaturated by fatty acid desaturases. (**A**) GC-FID quantification of gamma-linolenic acid 18:3n6 in H460 cells treated for 24 hours with BSA or 100 µM indicated fatty acid. (**B**) GC-FID quantification of arachidonic acid 20:4n6 in H460 cells treated for 24 hours with BSA or 100 µM indicated fatty acid. (**C**) RSL3 dose response for H460 cells pre-treated with 100 µM indicated fatty acid in the presence or absence of 10 µM SC-26196 FADS2 inhibitor for 24 hours. (**D**) Regioselectivity of mammalian fatty acid desaturases demonstrated on petroselaidic acid. (**E**) GC-FID quantification of oleic acid 18:1n9 in H460 cells treated for 24 hours with BSA or 100 µM indicated fatty acid. (**F**) GC-FID quantification of palmitoleic acid 16:1n7 in H460 cells treated for 24 hours with BSA or 100 µM indicated fatty acid. (**G**) RSL3 dose response for H460 cells pre-treated with 100 µM BSA in the presence or absence of 1 µM CAY10566 SCD inhibitor for 24 hours. (**H**) Lethal fraction of H460 cells pre-treated with 100 µM BSA in the presence or absence of 1 µM CAY10566 SCD inhibitor for 24 hours, then treated with 4 µM RSL3. (**I**) Immunoblot of H460 Tet-on (TO) SCD overexpression (OE) cell line following 24 hours of treatment with 2 µg/mL doxycycline (dox).

**Supplementary Figure S5.**
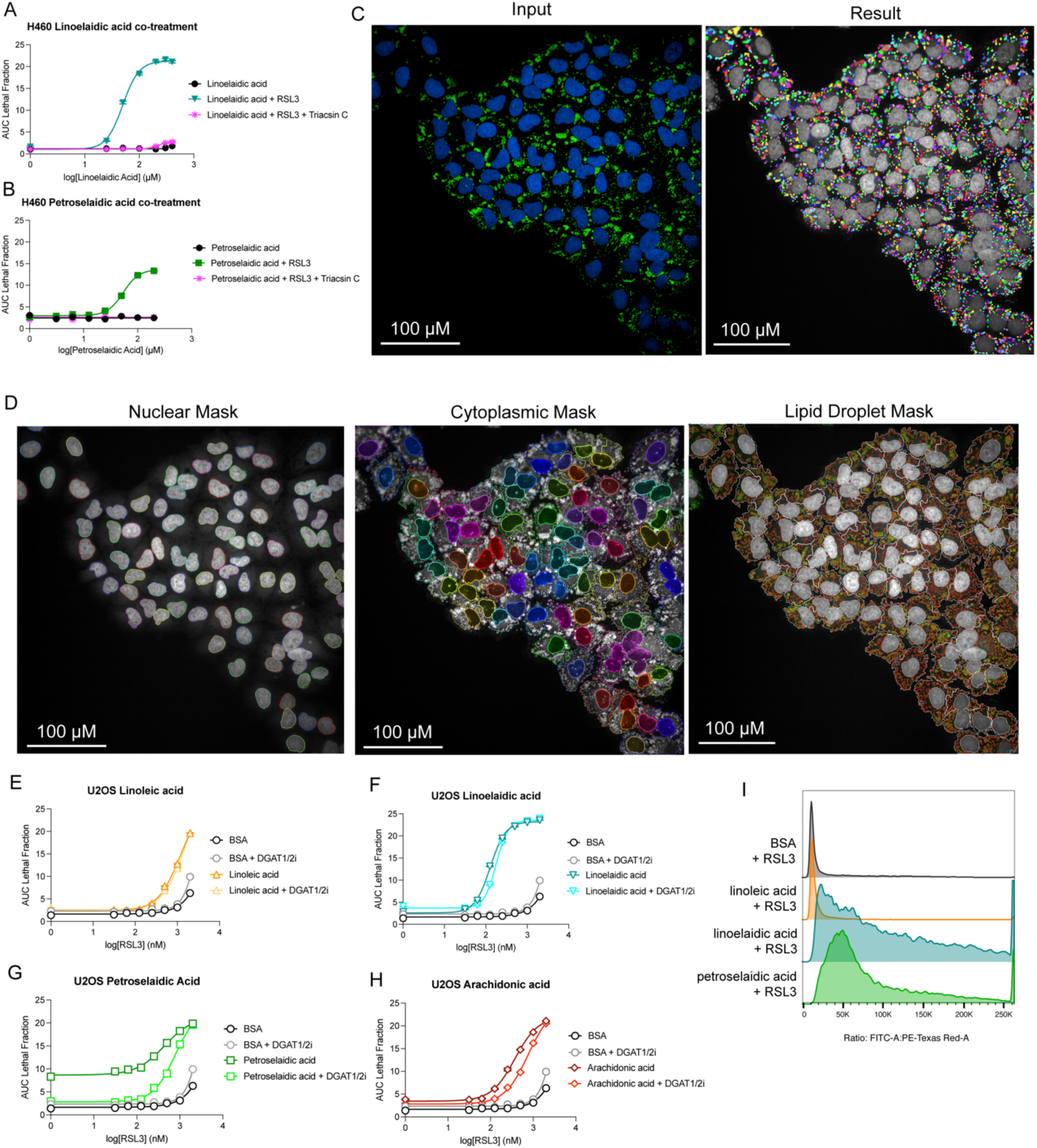
Exogenous fatty acid addition induces DGAT1/2-dependent lipid droplet formation with negligible effects on RSL3 dose response in U-2 OS. (**A**) Linoelaidic acid dose response of H460 cells co-treated with 1 µM RSL3 in the presence or absence of 5 µM Triacsin C. (**B**) Petroselaidic acid dose response of H460 cells co-treated with 1 µM RSL3 in the presence or absence of 5 µM Triacsin C. (**C**) Representative input and output images for lipid droplet quantification. Scale bar represents 100 microns. (**D**) Representative mask boundaries for nuclei, cytoplasm, and lipid droplets for lipid droplet quantification in H460 cells. Scale bar represents 100 microns. (**E**) RSL3 dose response of U-2 OS cells pre-treated with 100 µM linoleic acid in the presence or absence of 15 µM A-922500 DGAT1 inhibitor and 10 µM PF-06424439 DGAT2 inhibitor for 24 hours. (**F**) RSL3 dose response of U-2 OS cells pre-treated with 100 µM linoelaidic acid in the presence or absence of DGAT1 and DGAT2 inhibitors for 24 hours. (**G**) RSL3 dose response of U-2 OS cells pre-treated with 100 µM petroselaidic acid in the presence or absence of DGAT1 and DGAT2 inhibitors for 24 hours. (**H**) RSL3 dose response of U-2 OS cells pre-treated with 100 µM arachidonic acid in the presence or absence of DGAT1 and DGAT2 inhibitors for 24 hours. (**I**) Representative histogram of BODIPY C11 oxidized to non-oxidized fluorescence ratio in BSA or fatty acid pre-treated cells treated with RSL3 as quantified in Figure 5A.

